# Molecular Basis of Host-Adaptation Interactions between Influenza Virus Polymerase PB2 Subunit and ANP32A

**DOI:** 10.1101/2020.03.18.996579

**Authors:** Aldo Camacho Zarco, Sissy Kalayil, Damien Maurin, Nicola Salvi, Elise Delaforge, Sigrid Milles, Malene Ringkjøbing Jensen, Darren J. Hart, Stephen Cusack, Martin Blackledge

## Abstract

Avian influenza polymerase undergoes host adaptation in order to efficiently replicate in human cells. Adaptive mutants are localised on the C-terminal (627-NLS) domains of the PB2 subunit. In particular mutation of PB2 residue 627 from E to K in avian polymerase rescues activity in mammalian cells. A host transcription regulator ANP32A, comprising a long C-terminal intrinsically disordered domain (IDD), has also been shown to be responsible for this viral adaptation. Human ANP32A IDD lacks a 33 residue insertion compared to avian ANP32A, a deletion that restricts avian influenza polymerase activity in mammalian cells. We determined conformational descriptions of the highly dynamic complexes between 627E and 627K forms of the 627-NLS domains of PB2 and avian and human ANP32A. The negatively charged intrinsically disordered domain of human ANP32A transiently binds to a basic face of the 627 domain, exploiting multiple binding sites to maximize affinity for 627-NLS. This interaction also implicates residues 590 and 591 that are responsible for human-adaptation of the the 2009 pandemic influenza polymerase. The presence of 627E interrupts the polyvalency of the interaction, an effect that is compensated by extending the interaction surface and exploiting an avian-unique motif in the unfolded domain that interacts with the 627-NLS linker. In both cases the interaction favours the open, dislocated form of the 627-NLS domains. Importantly the two binding modes exploited by human- and avian-adapted PB2 are strongly abrogated in the cross interaction between avian polymerase and human ANP32A, suggesting that this molecular specificity may be related to species adaptation. The observed binding mode is maintained in the context of heterotrimeric influenza polymerase, placing ANP32A in the immediate vicinity of known host-adaptive PB2 mutants. This study provides a molecular framework for understanding the species-specific restriction of influenza polymerase by ANP32A and will inform the identification of new targets for influenza inhibition.

---

Influenza A virus (IAV) is responsible for 3-5 million severe cases every year, resulting in 250 to 500 000 deaths.^1^ Most influenza strains evolve exclusively in the large reservoir of water birds, but some highly pathogenic avian strains (e.g. H5N1, H5N8, H7N9) can infect humans with lethal consequences (up to 60% mortality) and are potential pandemic threats for humanity if they develop the human-to-human transmissability.^2^ However, for these avian *(av)* viruses to efficiently replicate in mammalian cells, host adaptation of the viral polymerase is necessary. Few mutations are required for this,^3–6^ and a number of these cluster on the surface of the C-terminal 221 amino acid section of PB2, comprising separate ‘627’ and ‘NLS’ domains.^4,7^ In particular mutation of residue 627 from E to K in *av*PB2 rescues polymerase activity and viral replication in mammalian cells. ^8–11^ Members of the host transcription regulator family ANP32A,^12^ comprising a long low-complexity acidic intrinsically disordered domain (IDD) at the C-terminus, have been shown to be responsible for this viral adaptation.^13^ *h*ANP32A lacks an insertion of 33 disordered residues compared to *av*ANP32A, restricting *av*H5N1 polymerase activity in mammalian cells (Figure 1A). This restriction is lifted by E627K mutation, suggesting an essential role for ANP32A through interaction with PB2,^14–22^ although there are currently no molecular descriptions of these interactions. The interaction between ANP32 and influenza polymerase is critical in supporting IAV replication and is attracting increasingly intense interest.^16–18^ Recent studies also point to the importance of related members of the ANP32 family, in particular ANP32B,^19–21^ as well as the role of surface residues in the folded leucine rich region (LRR) of ANP32A.^22^

**Figure 1.**
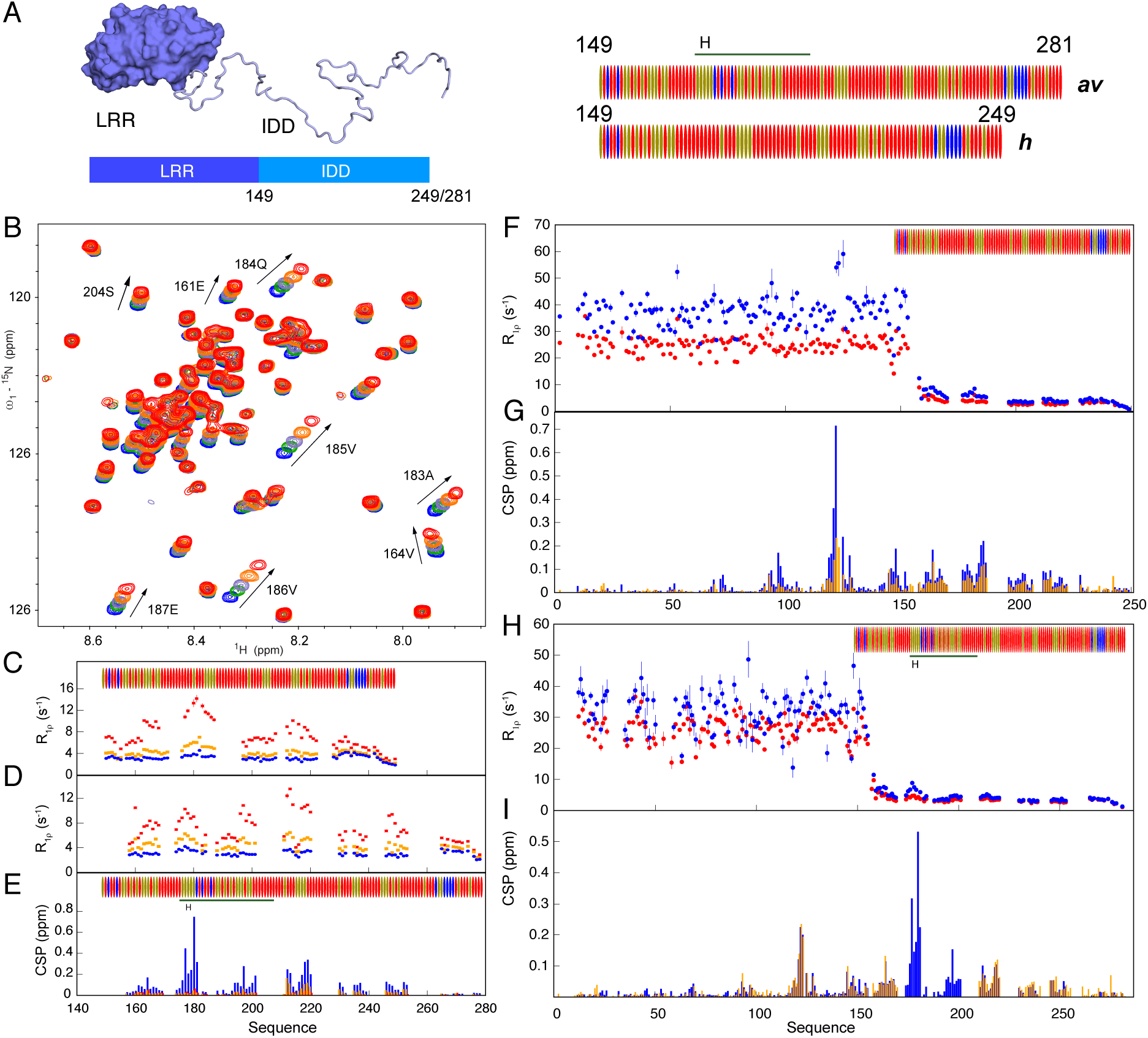
NMR of ANP32A with 627-NLS reveals a highly dynamic polyvalent complex. A. Representation of the domains of ANP32A. Blue surface shows the folded domain (leucine rich region – LRR), pale blue ribbon the disordered IDD that is 100 or 132 amino acids in length. The sequence distribution of the IDD is shown on the right. Red = Asp or Glu, blue = Arg or Lys, beige = hydrophobic or polar. The NLS sequence if visible as a basic patch at the C-terminus. The avian IDD is 32 amino acids longer, with a 33 amino acid inserted sequence (horizontal line) consisting of an avian-unique hexapeptide (H) followed by a 27 amino acid repeat sequence (182-188) replicating the sequence (149-175). B. Chemical shift titration of 627-NLS(K) into ^15^N labelled IDD of *h*ANP32A. *h*ANP32A was 25μM throughout. 627-NLS(K) concentrations were 0 (blue), 25 (green), 50 (grey-blue), 100 (orange) and 200 (red) μM. Measurement at 850MHz, 293K (see Methods). C. ^15^N rotating frame relaxation (R_1ρ_) of free (blue) ^15^N labelled IDD of *h*ANP32A at 300μM, and upon mixing with 627 (K) at 1:1 (orange) and 1:4 (red) ratio. Although the region around ^180^DEDA^183^ shows maximal increase in average relaxation rate, other sites up and downstream reveal additional interaction sites. Measurement at 850MHz, 293K. D. ^15^N rotating frame relaxation (R_1ρ_) of free (blue) ^15^N labelled IDD of *av*ANP32A at 300μM, and upon mixing with 627 (E) at 1:1 (orange) and 1:4 (red) ratio. Maximum effect is seen at the equivalent motif, 33 amino acids downstream from *h*ANP32A. E. Chemical shift perturbation (CSP) of full-length *av*ANP32A upon addition of 627(E) (orange), NLS (red) and 627-NLS(E) (blue). Concentration of *av*ANP32A was 25μM and the partner 50μM throughout. The strength of the CSP is significantly higher when both partners are present, substantiating indictations that the hexapeptide binds to the linker between 627 and NLS. Resonances from the LRR were not observable at this low concentration. F. ^15^N rotating frame relaxation (R_1ρ_) of free (red) ^15^N labelled full-length *h*ANP32A at 300μM, and upon mixing with 627-NLS(K) at 1:1 ratio (blue). Measurement at 850MHz, 293K. G. CSP of full-length *h*ANP32A upon addition of 627-NLS(K) (blue). Concentration of *h*ANP32A was 300μM and 627-NLS(K) 600μM. The cross interaction between *h*ANP32A and 627-NLS(E) is shown for comparison at the same stoichiometric ratio (orange). H. ^15^N rotating frame relaxation (R_1ρ_) of free (red) ^15^N labelled full-length *av*ANP32A at 300μM, and upon mixing with 627-NLS(E) at 1:1 ratio (blue). Measurement at 850MHz, 293K. I. CSP of full-length *av*ANP32A upon addition of 627-NLS(E) (blue). Concentration of *av*ANP32A was 300μM and 627-NLS(E) 600μM. The cross interaction between *h*ANP32A and 627-NLS(E) is shown for comparison at the same stoichiometric ratio (orange). The avian-specific insert is highlighted by shifting data from *h*ANP32A from residues following the beginning of the insert by 33 amino acids.

Conformationally, the 627-NLS region of PB2 exhibits intriguing behaviour.^7,23^ X-ray crystallographic structures of both *h* and *av-*adapted 627-NLS revealed a compact two-domain structure,^4,7^ a conformation also found in full-length transcriptionally-active polymerase.^24^ Solution NMR, however, revealed the coexistence of two forms of 627-NLS, corresponding to “open” and “closed” states that interchange in a highly dynamic equilibrium,^25^ while crystallographic investigation of the polymerase complex suggested a role for this open form in viral replication or polymerase assembly.^26,27^

Here we combine NMR and quantitative ensemble analysis to describe and compare the complexes formed between *av*ANP32A and a*v-*adapted 627-NLS (627-NLS(E)), and between *h*ANP32A and *h*-adapted 627-NLS (627-NLS(K)). Although the complexes are both found to be highly dynamic they exhibit significant differences. Polyvalent combinations of transient interactions between the acidic IDD and the positively charged 627K domain stabilize the *h*ANP32A:627-NLS(K) complex. This polyvalency is less efficient for *av*ANP32A:627-NLS(E), due to the interruption of an exposed basic surface on the 627 domain by the presence of E627. The weaker interaction is however compensated by the recruitment of additional sequences on the longer *av* IDD upon interaction with 627-NLS(E), in particular an avian-specific hexapeptide motif that interacts with the linker between 627 and NLS. Notably the cross-interaction between *h*ANP32A and 627-NLS(E) exhibits neither of these possible stabilisation mechanisms, which may explain the inability of IAV polymerase to function in human cells without the E627K mutation. Importantly, we show that the interaction exhibits the same properties in the presence of heterotrimeric influenza influenza polymerase, providing insight into the role of the IDD in the function of the ANP32A:polymerase complex.

## Investigation of the intrinsically disordered domains of h and avANP32A and their interactions with 627-NLS

We have compared the two complexes using solution state NMR, initially from the side of ANP32A. The intrinsically disordered domains (IDDs) of *h* and *av* ANP32A comprise 63/96 and 79/129 Asp or Glu residues respectively, leading to extensive spectral overlap. *h* and *av*ANP32A IDDs differ principally due to a 33 amino acid insert in *av*ANP32A (176-209) comprising an *av*-unique hexapeptide, ^176^VLSLVK^181^, followed by a duplication of 27 amino acids present in *h*ANP32A IDD. Backbone resonance assignment was completed to 78 and 58% respectively, revealing that *h* and *av*ANP32A IDDs are indeed both intrinsically disordered (figure S1, S2), with a slight tendency (20%) towards helical conformation for the hexapeptide.

Upon addition of the 627(K) domain, ^1^H and ^15^N chemical shift perturbations (CSPs) are seen for a large number of resonances in the IDD of *h*ANP32A (figures 1B, S1D). NMR relaxation rates measured at increasing titration admixtures (figure 1C) show maximal effects for the acidic strand ^180^DEDA^183^, while the largest CSPs are seen for the adjacent hydrophobic residues ^184^QVV^186^ (figure 1B, S1D). Additional interactions are seen throughout the chain, in particular at ^164^VE^165^ and ^214^YND^216^. Comparison of backbone ^13^C shifts in the free and fully bound states reveals that only ^179^YDED^182^ shows any evidence of folding upon binding (figure S1D), in this case into an extended *β*-sheet conformation, while the remainder of the chain remains highly flexible in the complex, retaining its random coil nature (figure S1B). Similar evidence of multiple interaction sites is seen for the IDD of *av*ANP32A in complex with the 627(E) domain (figure S1D and 1D). Relaxation properties (figure 1C, 1H) and more specifically CSPs (figure 1E) of the hexapeptide are strongly influenced by the presence of both domains, compared to constructs each comprising only the 627 or NLS domains and the linker. This implies that the specific conformational behaviour of the linker, or the relative position of the two domains, in integral 627-NLS is essential for the interaction with the hexapeptide of *av*ANP32A IDD.

Interaction of 627-NLS and ANP32A reveals similar profiles to those measured for the IDDs alone, with additional interactions involving the LRR for both *h* and *av*ANP32A (figures 1F-1I). In both *h* and *av*ANP32A, the largest CSPs in the LRR are centered on ^120^LFN^122^, with additional shifts induced following the spine of the beta-helix (figure 1G). Notably the cross interaction, between *h*ANP32A and 627-NLS(E) shows much smaller shifts than *h*ANP32A:627-NLS(K), while comparison with *av*ANP32A:627-NLS(E) clearly identifies increased shifts centered on the hexapeptide (figure 1I).

Although most of the IDD is involved in the interaction, the strongest binding, or highest populations of binding interactions, occur at a distance of 25-35 amino acids from the LRR. In the case of *av*ANP32A this concerns the avian-unique hydrophobic hexapeptide, and in *h*ANP32A the sequence ^180^DEDAQVV^186^. Further interactions are observed downstream of these interaction sites until the nuclear localization sequence (KRKR), situated approximately 15 amino acids from the end of the chain, beyond which point no significant interactions are observed.

Despite the evidence of clear interaction between ANP32A and 627-NLS, the complex is polyvalent, and in all cases weak (supporting information table S1, figure S3), with none of the individual interactions exhibiting a stronger affinity than 800μM for the interaction of *h*ANP32A with 627(K) (1700μM for *av*ANP32A with 627(E)). Notably the interactions between *h*ANP32A and 627(E) are also weaker (>1400μM), suggesting that the single mutation plays an important role. Interaction with integral 627-NLS is weaker due to the open-closed equilibrium reducing the population of available binding states. Interaction of the LRR of ANP32A alone with the 627 domain reveals weaker affinity (>1500μM). Although these interactions are weak, the extended interaction surface involving 80-100 disordered amino acids in *h* and *av*ANP32A nevertheless results in tighter binding as experienced by 627-NLS.

## Investigation of the interactions of avian and human adapted 627-NLS with h and avANP32A

The interaction was also investigated from the side of 627-NLS domains of PB2. The two domains exhibit an open-closed equilibrium (figure 2A) that is populated approximately 40:60 at 293K leading to two sets of resonances for the majority of the protein.^25^ Addition of *h*ANP32A to 627-NLS(K) resulted in CSPs throughout the protein (figure S4). The largest shifts are observed in the open form (figure S4), suggesting that ANP32A interacts preferentially with this conformation. The closed form of 627-NLS(K) is stabilized by a tripartite salt bridge, and can be removed from the equilibrium by mutation of the implicated amino acids (R650 or D730/E687).^25^ CSPs induced upon interaction with the ANP32A IDD are illustrated for clarity using an open-only mutant (figure 2B) and the distribution of CSPs as measured on the wild-type proteins (figure 2C). Figure 2D illustrates the major shifts observed for 627-NLS(K) upon addition of *h*ANP32A IDD. Although basic sidechains are found on both sides of the domain (figure 2E), CSPs are observed mainly on one face of the 627 domain, again suggesting that the interaction is not uniquely driven by electrostatic attraction with the acidic IDD.

**Figure 2.**
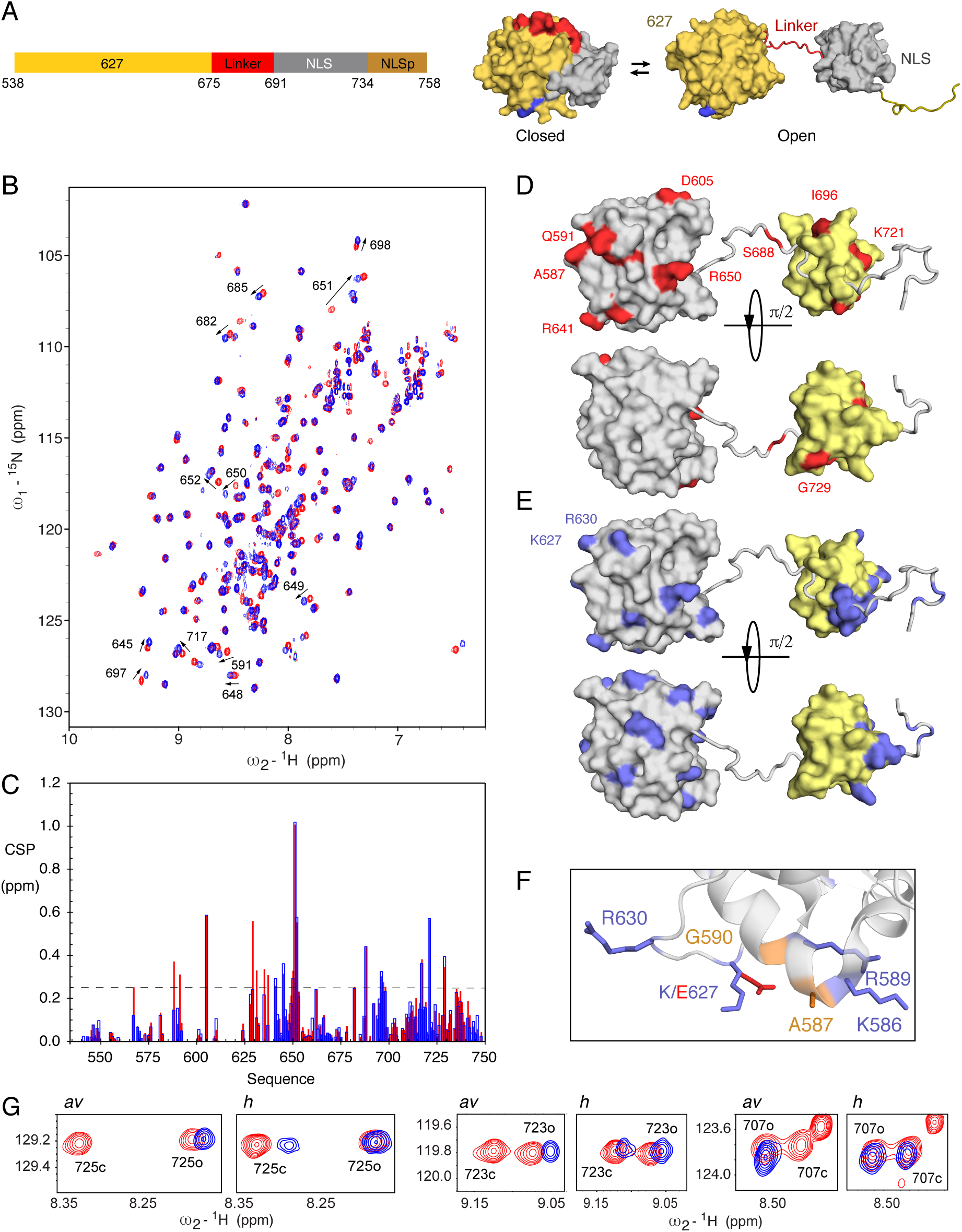
Interaction of 627-NLS and ANP32A observed from the perspective of 627-NLS. A. Representation of the domains of 627-NLS. The C-terminal domains of PB2 comprise two sub-domains (627 – orange and NLS - grey) connected by a linker (red) and terminated by an NLS peptide (NLSp). The molecule exists in equilibrium between closed and open forms that exchange at 50s^-1^ at room temperature. The position of the E627K adative mutation is shown in blue. B. HSQC of the open-only form of ^2^D, ^13^C, ^15^N labelled 627-NLS(K) (D730A/E687A) (200 μM) (red) indicating chemical shifts of selected sites upon addition of *h*ANP32A IDD (400 μM) (blue). C. CSP of 627-NLS(K) (300 μM) upon addition of *h*ANP32A IDD at a ratio of 1:4 (red) and 627-NLS(E) (300 μM) upon addition of *av*ANP32A IDD at a ratio of 1:4 (blue). Spectra recorded on ^2^D,^13^C, ^15^N labelled 627-NLS at 293K and 850MHz. Only shifts affecting resonances corresponding to the open form of the protein are shown for clarity. D. CSP of 627-NLS(K) (red) induced by addition of *h*ANP32A derived from (B) above the threshold of 0.25 (dashed line) for 627 (grey) and NLS (yellow) domains. Two orientations of the protein are shown. E. Distribution of basic sidechains (blue) on the surface of the 627 and NLS domains. By comparison with C it is evident that one of the two faces preferentially interacts with the IDD. F. The interaction of *h*ANP32A impacts the N-terminus of the 587-605 helix in 627-NLS(K), resulting in chemical shifts of residue 591. This region shows negligible CSP in the equivalent *av*ANP32A:627-NLS(K) interaction. Conformations taken from the structures 2vy7 and 2vy8.^4^ G. Interaction of full-length 627-NLS with ANP32A induces a change in the open-closed equilibrium. Red – peaks reporting on the open and closed forms of 627-NLS(E) or 627-NLS(K) and in the presence (blue) of *h* and *av*ANP32A and (blue) in the presence of 250μM of 627-NLS(K) and 627-NLS(E) respectively. The interaction with ANP32A potentiates the equilibrium in both cases, fully removing the closed form in the case of the avian pair. Residues distal from the main interaction site were chosen to reduce the risk of peak disappearance due to direct interaction.

Additional shifts are notable for 627-NLS(K):*h*ANP32A compared to 627-NLS(E):*av*ANP32A (figures 2C, S4), particularly in the vicinity of 587-591 and R630, indicating that the E627K mutation increases accessibility of the 587-591 region to the ANP32A IDD. It is interesting to note that residues 590 and 591 provide an alternative pathway to host adaptation, as evidenced in the 2009 pandemic strain.^5,28^ The long extended loop comprising 627E/K, that is highly flexible in both 627E and 627K (figure S5), wraps around the 588-605 helix, and interaction with *h*ANP32A appears to enhance the accessibility of the N-terminus of this helix uniquely when 627K is present (figure 2F).

The affinity measured when observing resonances from 627-NLS(K) is considerably tighter than from the side of *h*ANP32A, with values of 20μM for 627K, and 50μM for 627-NLS(K). This increased affinity apparently occurs due to avidity with the extensive interaction surface presented by *h*ANP32A. Although the stoichiometry cannot be determined accurately, titration curves imply that it is significantly different to 1:1 from the side of 627-NLS (figure S3), again suggesting that the increased affinity occurs due to polyvalent binding of multiple, weak binding sites on ANP32A to each site on the surface of 627-NLS(K). Notably the affinity for *av*ANP32A IDD measured when observing resonances of 627E is approximately 20 times weaker than for the equivalent human IDD:627K complex, with values >600μM. In combination with observations measured from the side of ANP32A, it therefore appears that the absence of 627K, which disrupts the continuity of the positively charged surface on 627, strongly abrogates this component of the interaction.

The interaction with ANP32A strongly favours the open form of 627-NLS (figure 2G), with the closed form essentially disappearing from the equilibrium at 1:2 ratio of 627-NLS(E):*av*ANP32A and falling from 65 to 40% at the same stoichiometry in 627(K)-NLS:*h*ANP32A.

## *Ensemble descriptions of the dynamic* 627-NLS(K):*h*ANP32A and 627-NLS(E):*av*ANP32A *complexes*

To develop a more detailed description of the dynamic interaction between 627-NLS and ANP32A, we have incorporated 8 cysteine mutants into both 627-NLS(K) and (E) and labelled the proteins with TEMPO-based paramagnetic spin label. This allows the detection of weak, or sparsely populated contacts between the two proteins, from the perspective of five positions on the 627 domain and three on the NLS domain (figure 3A-C). The experimental results show strong paramagnetic relaxation effects (PRE) distributed over long stretches of the IDD for certain spin-labels, and little broadening for other positions. The profiles again suggest polyvalent interaction, whereby distinct sites dispersed along the IDD visit the same sites on 627-NLS. Interestingly, two labels induce a well-defined pattern over the long *β*-sheet on the concave face of the LRR of ANP32A, allowing the determination of its orientation with respect to the 627 domain (see Methods). This orientation (figure S6) is in full agreement with the observed CSP on the surface of the LRR of ANP32A (figure 1), where the largest chemical shifts are seen for F121, Y122 and C123 which are positioned in closest proximity to the 627 domain. The interaction is apparently stabilized by hydrophobic interactions involving residues on the surface of 627 and ANP32A, and an electrostatic interaction between D130 (ANP32A) and R646 (627) (figure 3D). The former has recently been implicated in host adaptation.^22^ The optimal poses determined for the 627-NLS(K):*h*ANP32A and 627-NLS(E):*av*ANP32A complexes are essentially identical. The interface between 627 and ANP32A is bordered by a nearly continuous ridge of solvent accessible basic sidechains, including K589, R630, R641 and R650, that is completed by the presence of K627 in the case of 627-NLS(K) (figure 3E). Inspection of the PRE data from the IDD regions of *h* and *av*ANP32A reveals weaker effects in the region immediately following the LRR for *av*ANP32A, and more contacts with the dislocated NLS domain (figure 3A,B), in particular the spin-label at position 699 in the NLS broadens the IDD maximally at the position of the hexapeptide of *av*ANP32A. This again shows that the interaction with the immediately proximal basic face of 627 is weaker in the case of *av*ANP32A.

**Figure 3.**
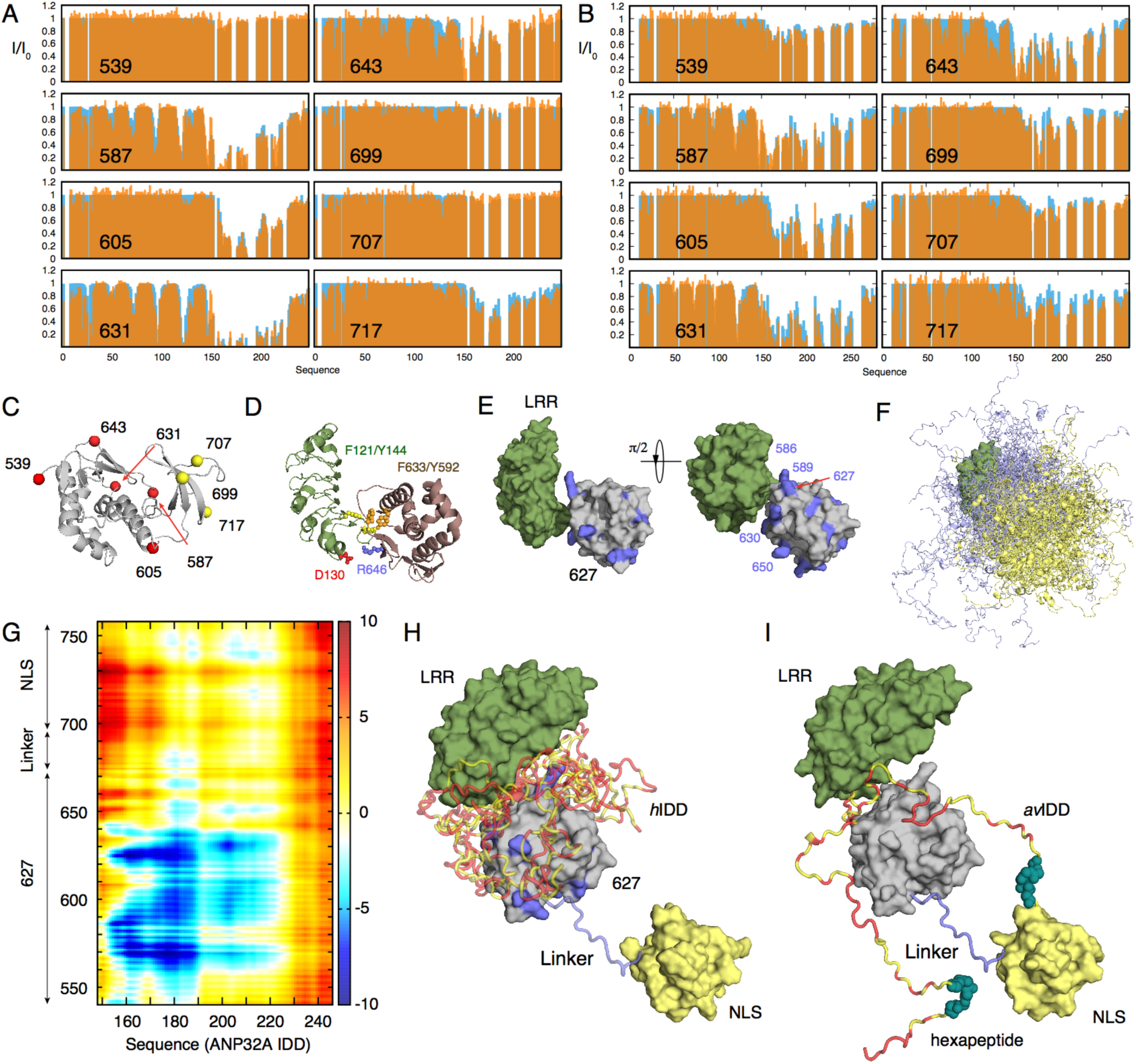
Characterisation of the dynamic 627-NLS(K):*h*ANP32A and 627-NLS(E):*av*ANP32A interaction complexes. A. Experimental (orange) and calculated (blue) paramagnetic relaxation enhancements measured on *h*ANP32A in the presence of paramagnetically labelled *h*ANP32A. Intensity ratios compare spectra recorded in the presence of oxidised and reduced forms of TEMPO-maleimide for complex admixtures of *h*ANP32A (300μM) and 627-NLS(K) (150μM). Calculated values result from representative ensembles selected using the ASTEROIDS approach. B. Experimental paramagnetic relaxation enhancements measured on *av*ANP32A in the presence of paramagnetically labelled *av*ANP32A. Intensity ratios compare spectra recorded in the presence of oxidised and reduced forms of TEMPO-maleimide for complex admixtures of *av*ANP32A (220μM) and *av*627-NLS (400μM). This admixture was chosen to replicate the population of bound state estimated in the case of 627-NLS(K):*h*ANP32A. C. Position of the eight cysteine mutations used to label 627-NLS. Five mutations were selected over the surface of the 627 domain and three on NLS. One mutant protein was expressed and purified for each site on both 627-NLS(K) and 627-NLS(E) and labelled with TEMPO-maleimide (see Methods). D. Localisation of intermolecular interactions possibly stabilising the interface between the two proteins, involving hydrophobic (F121 and Y144 in ANP32A and Y592 and F633 in 627-NLS), and electrostatic interactions (D130 in ANP32A and R646 in 627-NLS). E. Position of positively-charged ridge of solvent-exposed basic sidechains in the vicinity of the interface between the folded domains. F. Representative ensemble of conformations describing the conformational space sampled by the *h*ANP32A:627-NLS(K) complex. The LRR of ANP32A is shown in surface representation (green), the IDD is shown in light blue. 627 and NLS domains are shown in grey and light yellow cartoon representation. G. Average distance difference matrix, showing the average difference (*d*_*hh*_-*d*_*avav*_) in distance between amino acid positions in the IDD of ANP32A (x-axis) and the 627-NLS domains (y-axis) over the two ensembles. The colour code shown on the right is measured in Å. H. Representation of the key interactions between *h*ANP32A and 627-NLS(K). Polyvalent interactions between 627 and the IDD localise the disordered domain in the vicinity of the basic patch on the surface of 627 (see panel E). The different IDD chains are shown to represent different binding modes and are truncated at residue 200 for clarity. Red positions in the IDD indicate the acidic sidechains. Note that this representation (and panel I) illustrate the tendency over the entire ensemble that is highly disperse (see panel F and figure S7). I. Representation of the key interactions between *av*ANP32A and 627-NLS(E). In this case the average distance between the IDD and the surface of 627 is larger, but in general closer to the NLS domain, in particular the hexapaptide of ANP32A and the linker between 627 and NLS. The two IDD chains represent reduced polyvalency compared to *h*ANP32A and 627-NLS(K) and are again truncated at residue 200 for clarity.

Ensemble analysis of the PRE data using the ASTEROIDS approach,^29^ accounting for the flexibility in the 627-NLS linker and IDD domains of ANP32A, allows us to propose a molecular description of the conformational sampling of the entire complex. Representative ensembles of conformations of the multi-domain complex (figure 3F) reveal the sampling of conformational space of the IDD and NLS domains relative to the position of 627 and the LRR of ANP32A. Comparison of the position of the IDDs over the ensembles (figure S7) reveals more restricted sampling for *h*ANP32A, localising residues 175 to 200 in the vicinity of the basic ridge on the surface of 627. Sampling of *av*ANP32A is more dispersed (figure S7), as shown quantitatively in the average distance map (figure 3G), that identifies closer contacts in the 627-NLS(K):*h*ANP32A complex for residues 160, 180 and 205 with 570, 586 and 627, and closer contacts between the linker and NLS regions and *av*ANP32A region 150-200 (figure 3H, I).

Finally, comparison of PRE profiles of *h*ANP32A:627-NLS(E) demonstrates that this cross-interaction, that represents the case encountered when a non-adapted avian IAV infects human cells, shows less extensive and in general weaker contacts compared to *h*ANP32A:627-NLS(K) and *av*ANP32A:627-NLS(E) (figure S8). Absence of 627K diminishes the strong polyvalent contacts present in the former and the shorter IDD and lack of the hexapeptide abrogates the extensive interaction surface and the linker-specific contact present in the latter. The required components for interaction modes specific to either 627-NLS(K):*h*ANP32A or 627-NLS(E):*av*ANP32A are therefore both weakened when avian polymerase interacts with *h*ANP32A.

## Investigating the hANP32A:627-NLS complex in the context of full-length influenza polymerase

To determine whether the interactions characterized here are relevant in the context of a human-adapted full-length influenza polymerase we repeated the chemical shift titrations using the highly homologous 627-NLS domains from influenza B and heterotrimeric influenza B polymerase (bound to *v*RNA) with *h*ANP32A (figure 4A). The interaction sites are found to be the same as for 627-NLS from influenza A (figure 4B), and the first 50 amino acids of the IDD disappear from the spectrum (figure 4C), likely because the large particle results in extreme line-broadening for the residues most strongly involved in the interaction. It is interesting to speculate whether the interaction can be accommodated within known structures of the polymerase.^24,30^ Superposition of the 627 domain reveals that the LRR of ANP32A can be inserted into a broad cylindrical pocket formed by 627, the mid and cap-binding domains of PB2, also bordered by the C-terminal domain of PB1, allowing ANP32A to adopt the pose determined in solution (figure 4D). Interestingly the site of adaptive mutations 271^31^ in the mid-domain of PB2 and 521^32^ lie in the immediate vicinity of 627 and 590, while the recently identified 355^32^ faces the convex back surface of ANP32A on the opposite side of the pocket (figure S9), suggesting that the importance of these mutations involves interaction with ANP32A. Figures 4E and 4F illustrate the expected conformational space sampled by the IDDs of *av* and *h*ANP32A within the 627-NLS:ANP32A complexes, indicating a broader capture radius for the avian complex. A similar procedure was applied to the apo IAV polymerase,^33^ where the 627 domain sits on the surface, but is displaced relative to the polymerase core. In this case the 627-NLS:ANP32A complex would be easily accommodated (figure S9). The position of NLS on the surface of the polymerase, and its observed positional variability in existing structures, suggests that the open form of 627-NLS is sampled in both apo and RNA-bound polymerase.

**Figure 4.**
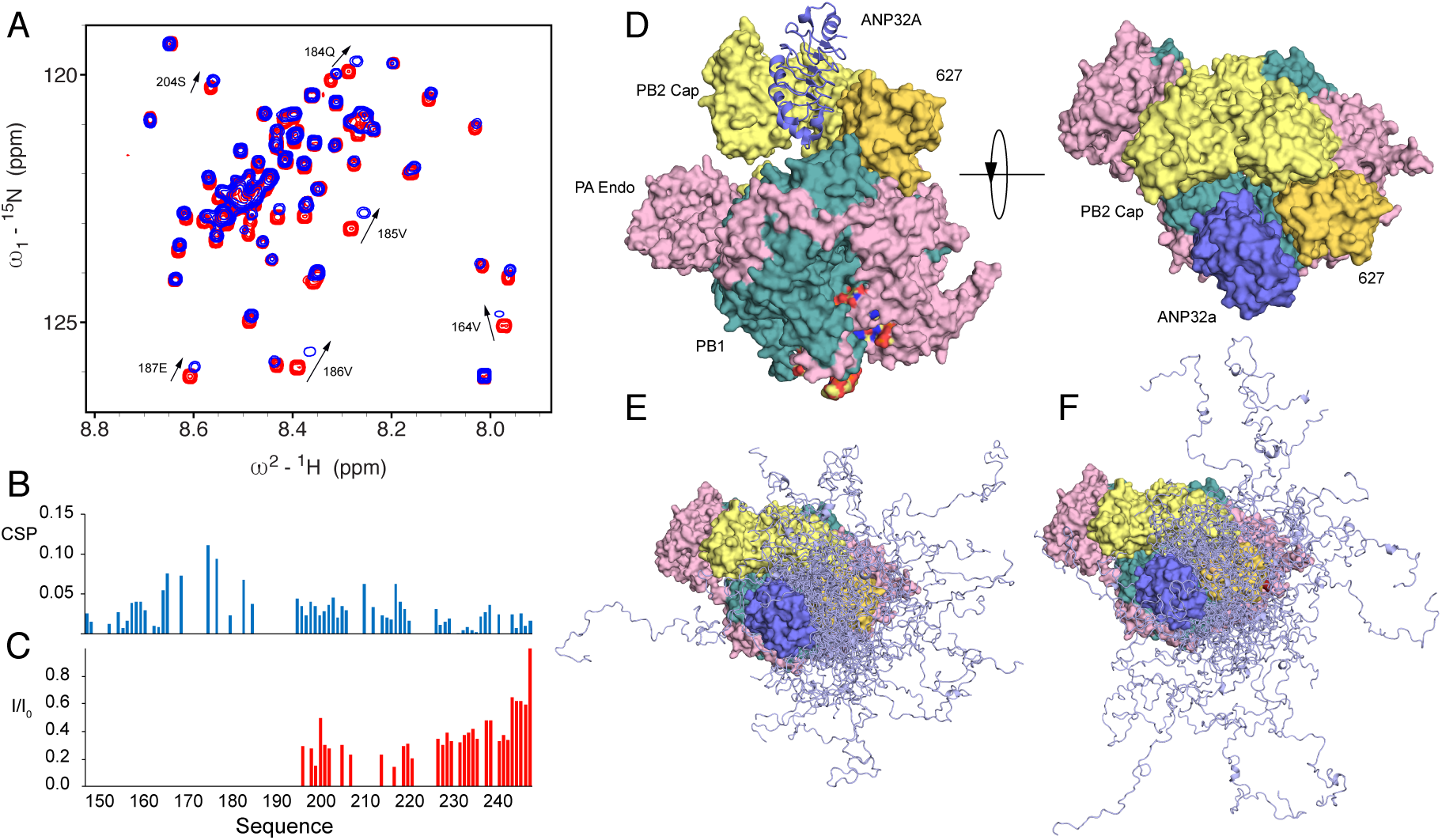
Interaction of hANP32A with full-length influenza B polymerase. A. CSP of ^15^N labelled IDD of *h*ANP32A (4μM) upon addition of full-length influenza B polymerase (32μM) bound to viral promoter RNA (*v*RNA). Measurement at 850MHz, 293K. Red - free protein, blue - in presence of polymerase. B. CSP associated with (A). C. Intensity ratio of ^15^N labelled full-length *h*ANP32A (4μM) upon addition of full-length influenza B polymerase (32μM) bound to *v*RNA. D. Compatibility of binding mode determined in free solution in the context of the vRNA bound conformation of influenza B.^24^ The 627 domain was superimposed on the 627 domain of PB2 in the full-length polymerase structure (4wsb). In this position, ANP32A LRR can be accommodated in a large pocket formed by 627 (yellow-orange) and cap binding domains of PB2 (yellow) and bordered by PB1 (dark-cyan). E. Conformational sampling of the IDD of *h*ANP32A, assuming the position of the LRR of ANP32A shown in D. The linker and NLS domains are not shown for clarity and are assumed flexible. F. Conformational sampling of the IDD of *av*ANP32A, assuming the position of the LRR of ANP32A shown in D. The linker and NLS domains are not shown for clarity and are assumed flexible.

## Discussion

In this study we describe and compare the molecular complexes formed by the human-adapted or avian-adapted 627-NLS domains with the respective ANP32A host proteins, in order to understand the nature and specificity of these interactions. All of the implicated proteins exhibit extensive intrinsic disorder. The elaboration of atomic resolution descriptions of such highly disordered complexes requires methodologies that can account for the ensemble of structures sampled by the two proteins in solution.^34^ Investigation of the *h*ANP32A:627-NLS(K) and *h*ANP32A:627-NLS(E) complexes using NMR chemical shifts, spin relaxation and paramagnetic relaxation combined with quantitative ensemble modelling reveals the existence of highly dynamic molecular assemblies that exhibit very different interaction modes.

The LRR of ANP32A interfaces with the 627 domain of PB2 at the C-terminal end of its concave surface, apparently stabilized via hydrophobic and electrostatic interactions. The disordered domain interacts transiently with a basic patch on the surface of the 627 domain, most strongly in the case of *h*ANP32A:627-NLS(K), involving weak, polyvalent interactions of acidic and hydrophobic residues of the IDD. The presence of multiple low-affinity interactions (in the millimolar range individually) distributed throughout 80 amino acids of the IDD of *h*ANP32A results in an effective increase in affinity to around 50μM for 627-NLS(K), an effect of avidity that has been observed in a number of systems exhibiting extensive intrinsic disorder.^35,36^ The critical E627K mutation completes a continuous ridge of solvent-exposed positively charged residues that are available for interaction with the highly dynamic acidic IDD.^4^ This localizes interacting residues of the IDD in the vicinity of this surface, also implicating residues between 589 and 591 on 627-NLS only when the E627K mutation is present. Mutation of these sites was implicated in the establishment of the 2009 pandemic strain of IAV, underlining the possible importance of this polyvalent interaction in host adaptation.

By contrast *av*ANP32A:627-NLS(E) populates fewer conformations in the vicinity of the surface of 627. This lack of interaction is compensated by an even broader conformational sampling of the IDD that exploits a longer interaction surface, implicating the hexapeptide motif specific to *av*ANP32A and the NLS domain and in particular the linker region of 627-NLS. It is again interesting to note that two adaptative-mutations (V683T and A684S)^31^ have been identified in this linker region. By binding predominantly to the open form of 627-NLS, ANP32A potentiates the equilibrium of open and closed forms of 627-NLS, an effect that is more efficient in the case 627-NLS(E):*av*ANP32A interaction, likely as a result of the interaction of the hexapeptide. Notably *h*ANP32A:627-NLS(E) exhibits neither the stabilization properties mediated by the *av*IDD in *av*ANP32A:627-NLS(E) or the polyvalent binding specific to 627K as observed in the *h*ANP32A:627-NLS(K), resulting in fewer and weaker contacts. These effects may explain the inefficiency of avian polymerase in human cells in the absence of *av*ANP32A or 627-NLS(K).

Our results indicate that the complex characterized for the minimal 627-NLS:ANP32A interaction is maintained in the context of the integral polymerase. Interestingly the relative position of 627 and the LRR of ANP32A would place the host protein in a large pocket bounded by PB2 domains. Intriguingly, in this conformation the host-adaptive PB2 mutants lie in the immediate vicinity of ANP32A. The proximity of ANP32A IDD to the interface between the 627 and mid-domains of PB2, both of which undergo large-scale reorientations and dislocation between apo- and *v*RNA-bound polymerase, also raises the possibility that ANP32A interaction is associated with these conformational changes. The apo-polymerase conformation is also compatible with the binding pose determined here.

Recent observations have established that dimerization of influenza polymerase is essential for the initiation of *v*RNA synthesis during replication.^33^ It has been suggested that ANP32A plays a role in assembly or regulation of this dimerization process, for example, by recruitment of a second polymerase to apo polymerase for the formation of viral RNPs.^37^ In this context the polyvalent nature of the interaction between ANP32A and 627-NLS may be of functional relevance. In particular, the more extensive effective capture radius of the IDD of *av*ANP32A may be important.

In summary, the description of these highly dynamic assemblies reveals unique mechanistic insight into the role of the ANP32 family in host adaptation of avian influenza polymerase to the human cells and provides a molecular framework for understanding the considerable volume of experimental observation measured on this complex system as well as informing the identification of novel targets for IAV inhibition.

## Data Availability

All of the NMR data presented in the article are available from the authors upon request. The NMR chemical shift assignments of avian and human ANP32A will be deposited in the Biological Magnetic Resonance Bank (BMRB).

## Reporting Summary

A Nature Research Reporting Summary is deposited with this article describing research design and materials.

## Additional Information

Supporting information is available for this paper. Correspondence and requests for materials should be addressed to MB

## Acknowledgements

This project has received funding from the European Union’s Horizon 2020 research and innovation programme under the Marie Skłodowska-Curie grant agreement No. 796490 (AvInfluenza) and HFSP postdoctoral HFSP fellowship LT001544/2017 *“Molecular basis of avian influenza polymerase adaptation to human hosts”*. This work used the platforms of the Grenoble Instruct-ERIC center (ISBG; UMS 3518 CNRS-CEA-UGA-EMBL) within the Grenoble Partnership for Structural Biology (PSB), supported by FRISBI (ANR-10-INBS-05-02) and GRAL, financed within the University Grenoble Alpes graduate school (Ecoles Universitaires de Recherche) CBH-EUR-GS (ANR-17-EURE-0003). IBS acknowledges integration into the Interdisciplinary Research Institute of Grenoble (IRIG CEA).

## Author Contributions

ACZ and MB designed experiments. DM, ED, SK and ACZ prepared proteins. ACZ carried out NMR experiments. MB, SM, NS and MRJ supervised experimental work. ACZ and MB analysed and interpreted experimental data in terms of molecular complexes. SC provided reagents. SC and DH discussed interpretation with ACZ and MB. MB wrote the manuscript, and incorporated comments fro all authors.

## Author Contributions

Correspondence to Martin Blackledge.

## Ethics declaration

The authors have no competing interests.

## Methods

### Constructs

A codon optimized 627-NLS construct from PB2 subunit was synthesized encoding aa 538-759 from avian H5N1 A/duck/Shantou/4610/2003 for expression in *E. coli* (Geneart, Regensburg, Germany) as described.^25^ In addition, constructs containing just the 627 domain (aa 538-693) or the NLS domain (aa 678-759) were generated. Avian ANP32A (*Gallus gallus*, XP_413932.3) was synthesized and codon optimized for expression in *E. coli* (GenScript, New Jersey, USA). Plasmids containing just the intrinsically disordered region of avian ANP32A (avIDD, aa 149-281) or human ANP32A (hIDD, aa 144-249) were generated. All constructs were cloned into a plasmid derived from pET9a with an N-terminal His-tag and a TEV cleavage site (MGHHHHHHDYDIPTTENLYFQG). pQTEV-ANP32A was a gift from Konrad Buessow (Addgene plasmid # 31563; http://n2t.net/addgene:31563; RRID:Addgene_31563).^38^

### Protein expression and purification

Plasmids were transformed into *E. coli* Rosetta cells and the cultures were grown in LB and induced with IPTG for 16 hours at 18 °C. Bacteria were harvested by centrifugation, resuspended in buffer A (50 mM Tris-HCl pH 7.5 and 200 mM NaCl) with protease inhibitors (complete, Roche) and bacterial lysis was performed by sonication. All proteins were purified by affinity chromatography on Ni-NTA agarose (Qiagen), followed by incubation with TEV protease at 4 °C coupled with dialysis into buffer A. A second affinity column with Ni-NTA agarose was performed and the flow-through was loaded into a Superdex 75 column (GE Healthcare) for size exclusion chromatography in buffer A.

To produce ^15^N-labelled or ^15^N and ^13^C-labelled proteins for NMR spectroscopy, bacteria were grown in M9 minimal medium containing MEM vitamins (Gibco), supplemented with 1g/L of ^15^NH_4_Cl and 2 g/L of unlabeled or ^13^C-glucose. To produce additionally ^2^H-labelled proteins, the M9 minimal medium was prepared in D_2_O and 2 g/L of deuterated ^13^C-glucose. Protein purity was checked by SDS-PAGE and mass spectrometry. Single point mutations of 627-NLS were done using the Quick change method,^39^ using Phusion high fidelity DNA polymerase and DpnI (Thermo Scientific). Cysteine mutants were purified as mentioned above for wild-type protein, however 10 mM of dithiothreitol (DTT) was added after the second Ni-NTA column to keep proteins in a reduced state until labeling. Full-length human influenza polymerase from B/Memphis/13/03 (FluB) was overexpressed and purified as reported.^30^

### NMR spectroscopy

All samples for NMR were measured in 50 mM Tris-HCl buffer pH 6.5, 200 mM NaCl and 10% D_2_O. The assignment of the intrinsically disordered regions of avian and human ANP32A were obtained using ^15^N,^13^C-labeled samples (700μM) using BEST-TROSY tridimensional experiments recorded on a Bruker spectrometer equipped with a cryoprobe operating at 20 °C and a ^1^H frequency of 850 MHz. All spectra were processed using NMRPipe^40^ and analyzed in Sparky.^41^ MARS^42^ was used for spin system identification, followed by manual verification. The folded domain of ANP32A has been assigned previously.^43,44 13^C*α* chemical shifts of the intrinsically disordered regions were compared to random coil values using the software SSP.^45^

^15^N R_1_*p* relaxation rates were measured at 293K and a ^1^H frequency of 850 MHz using a spin lock of 1.5 kHz as described.^46^ A typical set of relaxation delays included points measured at 1, 15, 30, 50, 100, 140, 200 and 230 ms, including repetition of one delay. Relaxation rates were determined using in-house software and errors were estimated on the basis of noise-based Monte Carlo simulation. Interaction experiments with full-length polymerase were acquired with ^15^N-labelled hIDD or full-length ^15^N *h*ANP32A at a concentration of 4 μM after the addition of 32 μM of human FluB polymerase bound to the 5’ end viral RNA promoter (5’-pAGUAGUAACAAGAG-3’ OH). These experiments were recorded at 293K and a ^1^H frequency of 850 MHz.

PRE effects used to model the complex formed by ANP32A (^15^N-labeled, human or avian) and 627-NLS (627E or 627K) were measured from the peak intensity ratios between a ^15^N-HSQC 2D spectrum recorded on a sample containing 627-NLS labelled with TEMPO and a reference diamagnetic sample which was incubated previously with 5 mM of DTT. For these experiments, single cysteine mutants at positions 539, 587, 605, 631, 643, 699, 707 and 717 were tagged using 4-Maleimido-TEMPO. Briefly, purified 627-NLS single cysteine mutants were reduced with 10 mM of DTT at 4 °C for 12 hours and then dialysed throughly into 50 mM phosphate buffer pH 7.0 containing 150 mM NaCl without DTT. A 5-fold molar excess of 4-Maleimido-TEMPO dissolved in DMSO was added to the reduced 627-NLS cysteine mutants. The reaction was incubated for 12 hours at 4 °C and then injected into a Superdex S75 column to eliminate the excess of TEMPO through size exclusion chromatography. Complete labeling with TEMPO was verified by mass spectrometry. Measurement of PRE effects was done in samples containing 200 mM of ^15^N-labeled hIDD and 100 or 200 mM of the respective TEMPO-labeled 627-NLS (627K) mutants. Measurements in full-length *h*ANP32A were done with 300 mM of ^15^N-*h*ANP32A and 150 mM of the 627-NLS (627K) mutants and measurements on full-length *av*ANP32A were done with 220 mM of ^15^N labelled protein and 400 mM of the 627-NLS (627E) mutants.

### Determination of the relative position of ANP32A and 627

Experimentally determined paramagnetic relaxation enhancements (PREs) measured on *h* and *av*ANP32A in the presence of different spin-labelled forms of *h* and *av*627-NLS respectively were used to determine the relative position of the folded domain of ANP32A with respect to 627. Two tousand different positions of the two domains were generated using the program Haddock, varying over a wide range of distances and orientations. Positions of spin-label bearing sidechains were generated on the basis of rotameric libraries as described previously^29,47^ (figure S6) and an ensemble of sidechain positions was used to calculate expected PREs on ANP32A for a given position of 627. Admixtures were adjusted to ensure a population of the bound state of 10% for both complexes. The position of each of the 2000 starting conformations were varied over a range of ±10Å along three orthogonal cartesian axes at a resolution of 0.1Å, and the best fitting position retained. The ten best fitting structures are shown in figure 3D.

### Ensemble description of *h*ANP32A: 627-NLS(K) and *av*ANP32A:627-NLS(E) complexes

Having determined the relative position of the two domains, the flexible parts were constructed onto this conformation. For both *h* and *av* complexes, the statistical coil model flexible meccano^48^ was used to predict 10000 conformations of the linker region of 627-NLS, the NLS domain, the NLS peptide that terminates 627-NLS and the 96 or 128 amino acid IDD of *h* and *av*ANP32A. Conformers were calculated using amino acid specific potentials that reproduce the experimentally observed behaviour of the IDD domains, and were calculated to avoid steric overlap between any of the domains (figure S6). PREs over the entire ANP32A molecule (folded and unfolded domains) were calculated for each of the conformers calculated for each complex and these conformations were used as a basis set from which ensembles were selected using the ASTEROIDS approach.^34,49^ Ensemble size was estimated on the basis of direct and cross-validated PRE profiles (60 conformers were used for both *h* and *av* complexes).

Distance matrices were calculated by calculating the average distance between C*α* atoms between the two proteins in the selected ensembles from the two complexes, and the distance-difference matrix shown in figure 3F by subtracting the matrix from *h*ANP32A:627-NLS(K) from the *av*ANP32A:627-NLS(E) matrix.

## Supporting Information

**Table S1.** Affinities of different constants. A – All K_D_s were estimated using NMR spectroscopy via chemical shift titrations (see figure S5)

| Residue | Kd ($\mu\text{M}$)<sup>a</sup> | Std Error |
| --- | --- | --- |
|  | **<sup>15</sup>N -627E + avIDD** |  |
| 651 | 630.58 | 218.8 |
| 652 | 743.08 | 257.6 |
| 662 | 994.59 | 229.4 |
| 682 | 871.17 | 224.2 |
|  | **<sup>15</sup>N -627K + hIDD** |  |
| 588 | 16.22 | 1.99 |
| 591 | 20.19 | 4.04 |
| 645 | 17.57 | 3.16 |
| 651 | 20.22 | 2.99 |
| 652 | 30.35 | 7.44 |
| 681 | 34.3 | 12.7 |
| 682 | 36.74 | 12.06 |
|  | **<sup>15</sup>N -NLS + avIDD** |  |
| 688 | 457.63 | 11.9 |
| 690 | 430.8 | 25.6 |
| 738 | 522.3 | 41.8 |
| 739 | 400.82 | 39.15 |
|  | **<sup>15</sup>N -NLS + hIDD** |  |
| 688 | 605.04 | 55.63 |
| 690 | 602.36 | 23.19 |
| 738 | 583.44 | 50.56 |
| 739 | 547.73 | 84.14 |
|  | **<sup>15</sup>N -hIDD + 627E** |  |
| 178 | 2350.4 | 641.7 |
| 183 | 1699.9 | 191.2 |
| 184 | 1436.6 | 230.3 |
| 185 | 1898.5 | 109 |
| 186 | 1797.2 | 123.1 |
|  | **<sup>15</sup>N -hIDD + 627K** |  |
| 183 | 779.3 | 91 |
| 184 | 817.1 | 81.4 |
| 185 | 965.3 | 88.7 |
| 186 | 1008.3 | 59.1 |
| 187 | 867.1 | 104.9 |
|  | **<sup>15</sup>N -*av*IDD + 627E** |  |
| 217 | 1730.2 | 335.4 |
| 218 | 1884.5 | 294.2 |
| 219 | 1807.4 | 314.7 |
| 220 | 1848.0 | 274.3 |
|  | **<sup>15</sup>N -*h*LRR + 627-NLS (K)** |  |
| 119 | 1575 | 82.9 |
| 120 | 1700 | 100.2 |
| 121 | 1670 | 58 |
| 122 | 1659 | 61 |
|  | **<sup>2</sup>H, <sup>15</sup>N -*h*ANP32A + 627NLS(K)** |  |
| 98 | 1581 | 59 |
| 122 | 1407 | 79.7 |
|  | **<sup>2</sup>H, <sup>15</sup>N -*av*ANP32A + 627NLS(E)** |  |
| 122 | >3000 |  |
|  | **<sup>2</sup>H, <sup>15</sup>N-627NLS(K) + *h*ANP32A** |  |
| 644o | 28.9 | 6.44 |
| 645o | 42.2 | 3.6 |

**Figure S1.**
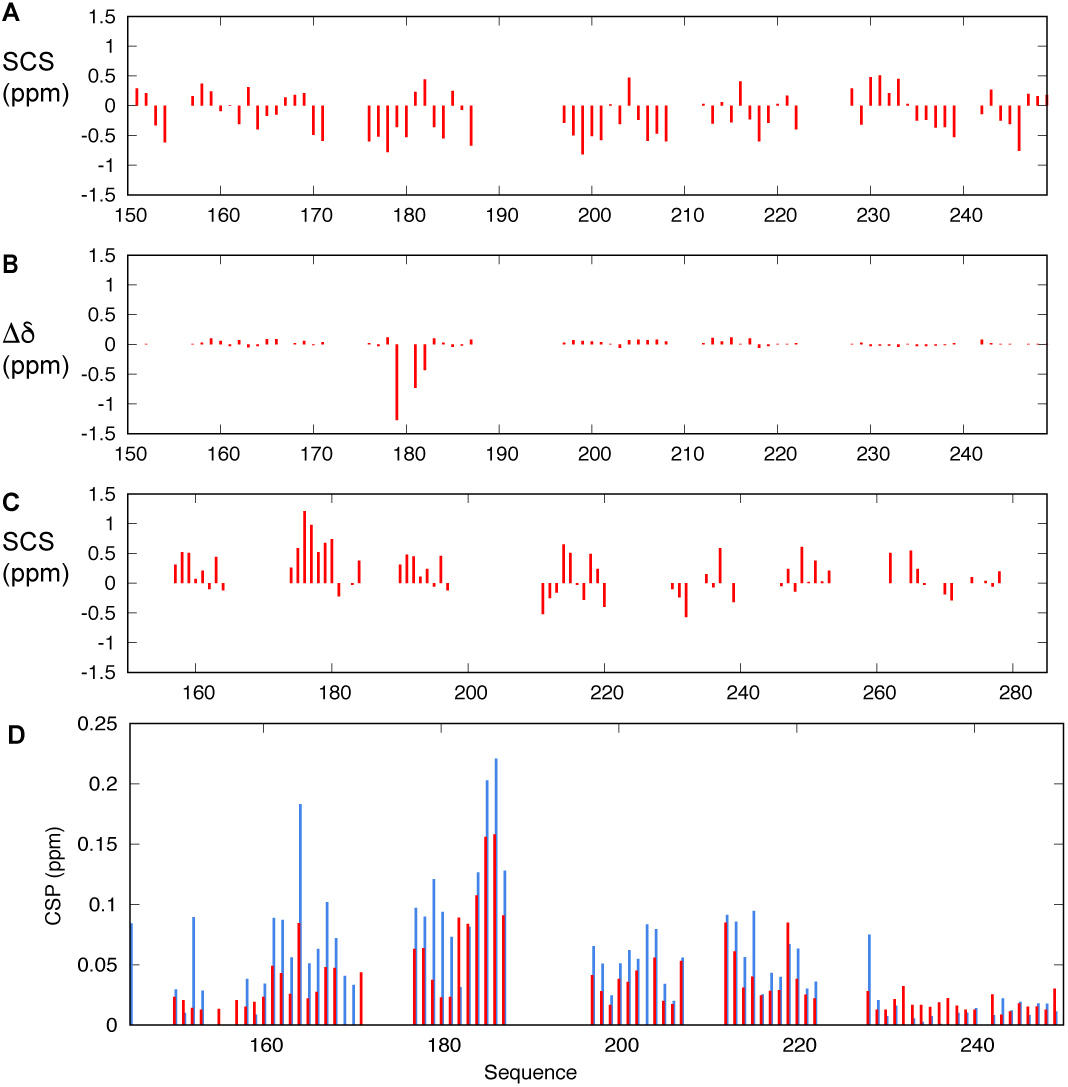
ANP32a C-terminal domains adopt intrinsically disordered conformations in solution. A. Secondary ^13^C*α* chemical shifts along the sequence of *h*ANP32a IDD. Values are all close to zero, indicating negligible propensity for secondary structure in solution. B. ^13^C*α* chemical shift perturbation (CSP) of *h*ANP32a IDD upon interaction with *h*627-NLS. Chemical shifts were determined from heteronuclear 3D BEST-TROSY three dimensional experiments. Only the ^179^YDED^182^ region shows evidence of conformational change upon binding. C. Secondary ^13^C*α* chemical shifts along the sequence of *av*ANP32a IDD. Values are again close to zero, indicating negligible propensity for secondary structure in solution except for the hexapeptide ^176^VLSLVK^181^ that shows weak (approximately 20%) helical propensity. D. Comparison of ^15^N, ^1^H chemical shift perturbation between *h*ANP32a IDD:627K (red bars) and *h*ANP32a:*h*627-NLS (blue bars). *h*ANP32a IDD:627K spectra were recorded at 293K with concentrations of 300 μM (1:1 mixture). *h*ANP32a: *h*627-NLS spectra were recorded at 293K, with concentrations of 25 and 50μM respectively.

**Figure S2.**
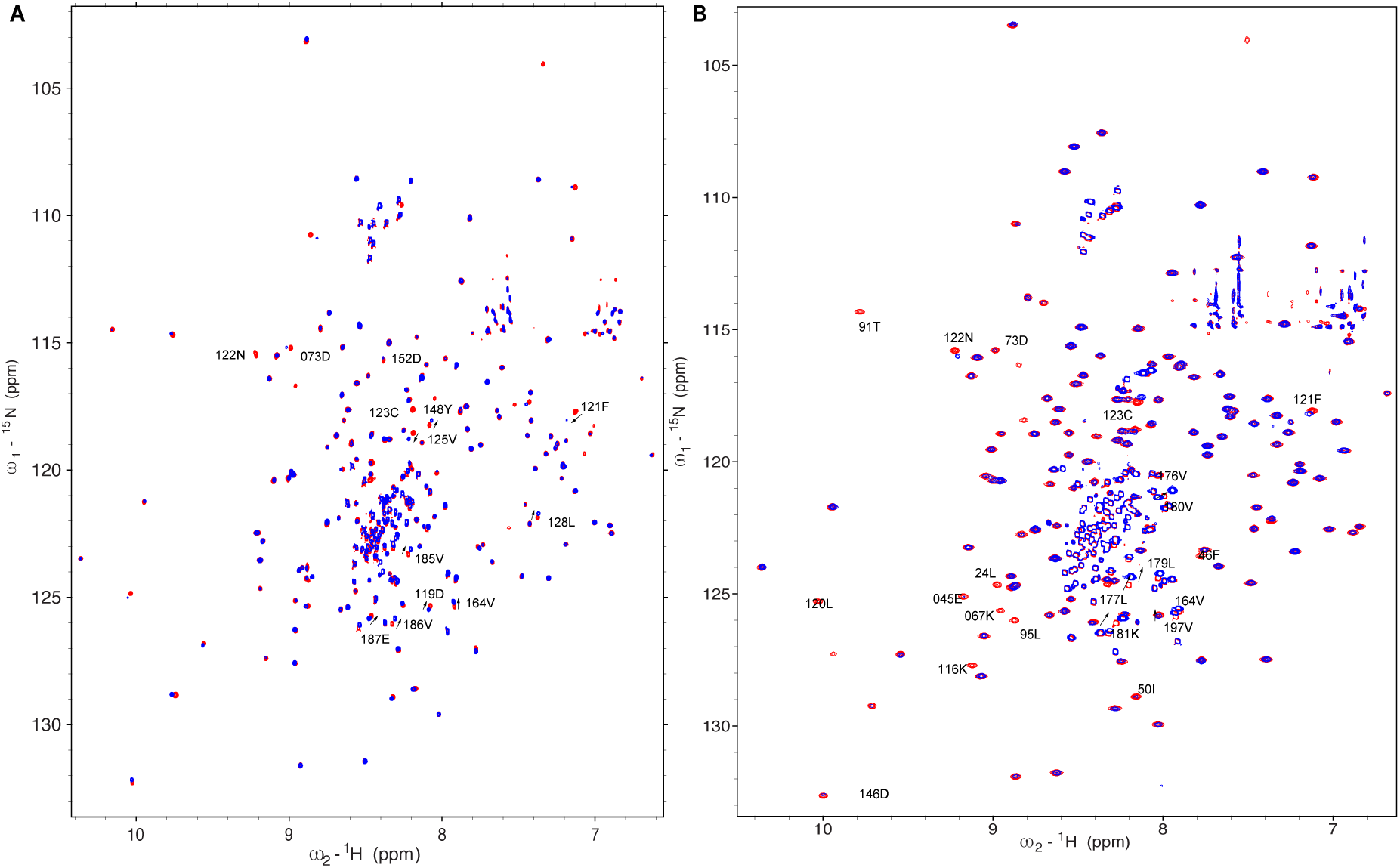
A. HSQC of (red) ^15^N labelled *av*ANP32a (300μM) upon addition of *av*627-NLS (600μM) (blue). Data were recorded at 950MHz, 293K. B. TROSY of (red) ^15^N labelled *av*ANP32a (300μM) upon addition of *av*627-NLS (600μM) (blue). Data were recorded at 700MHz, 293K.

**Figure S3.**
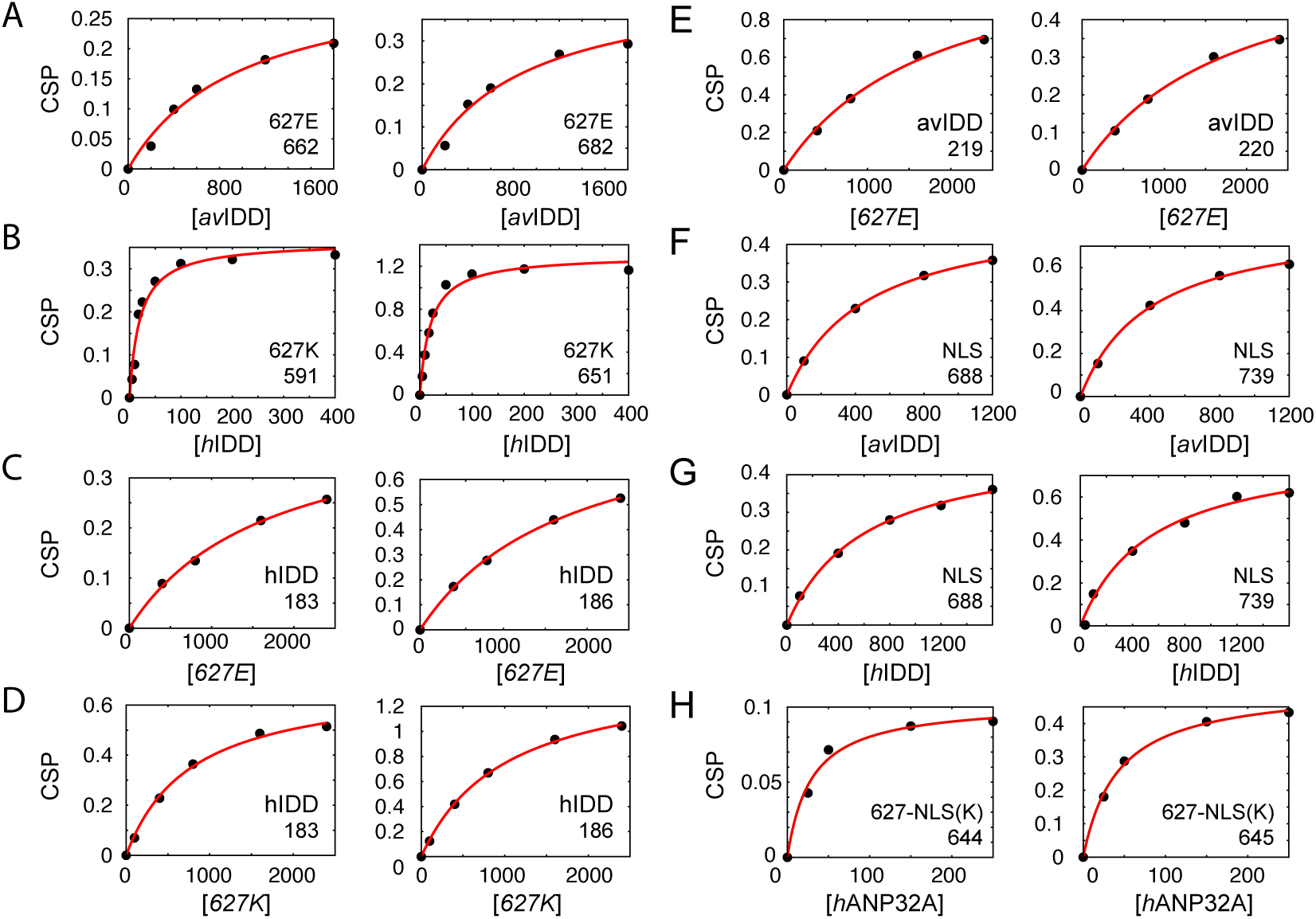
Examples of chemical shift titrations associated with K_D_ values estimated in table S1. Concentrations of the titrated partner are shown on the x-axis in μM. The residue number is given for each peak titration. Concentrations of the observed, labelled proteins were: A) 200 μM of ^15^N-627E, B) 200 μM of ^15^N,^13^C-627K, C) 200 μM of ^15^N-hIDD, D) 200 μM of ^15^N-hIDD, E) 200 μM of ^15^N,^13^C- avIDD, F) 180 μM of ^15^N-NLS, G) 180 μM of ^15^N-NLS and H) 250 μM of ^2^H,^15^N-627-NLS(K).

**Figure S4.**
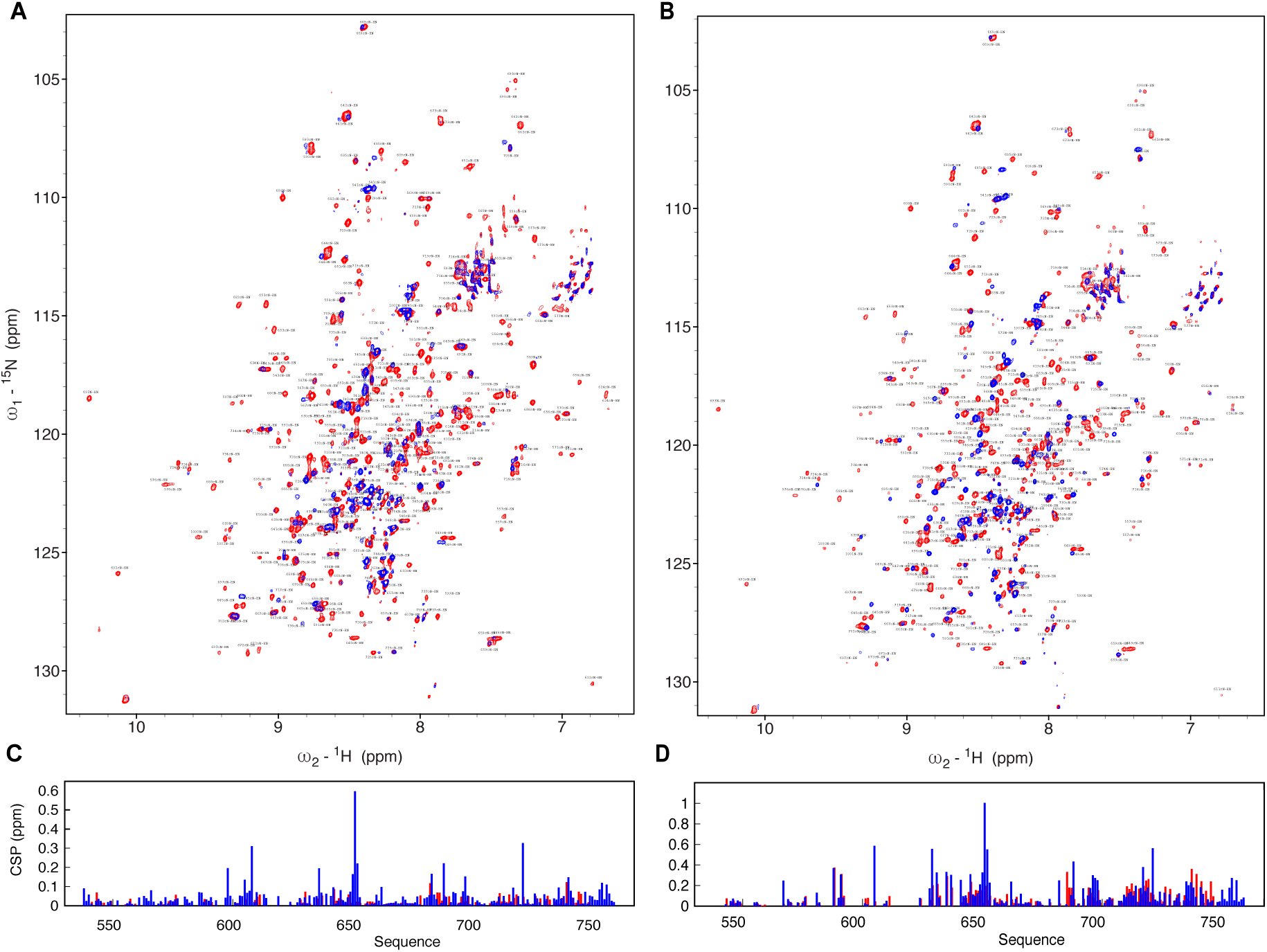
A. TROSY spectra of ^2^D, ^15^N labelled *h*627-NLS alone (red) (250μM) and in the presence of unlabelled of *h*ANP32a (250μM). Spectra acquired at 950MHz and 293K. B. TROSY spectra of ^2^D, ^15^N labelled *av*627-NLS alone (red) (250μM) and in the presence of unlabelled of *av*ANP32a (250μM). Spectra acquired at 950MHz and 293K. C. Chemical shift perturbation (CSP) of *av*627-NLS (250 μM) upon addition of full length *av*ANP32a at a ratio of 1:1. Blue: resonances corresponding to the open form. Red: resonances corresponding to the closed form. D. Chemical shift perturbation (CSP) of *h*627-NLS (250 μM) upon addition of full length *h*ANP32a at a ratio of 1:1. Blue: resonances corresponding to the open form. Red: resonances corresponding to the closed form. All spectra were recorded on ^2^D, ^13^C, ^15^N labelled 627-NLS at 293K and 850MHz.

**Figure S5.**
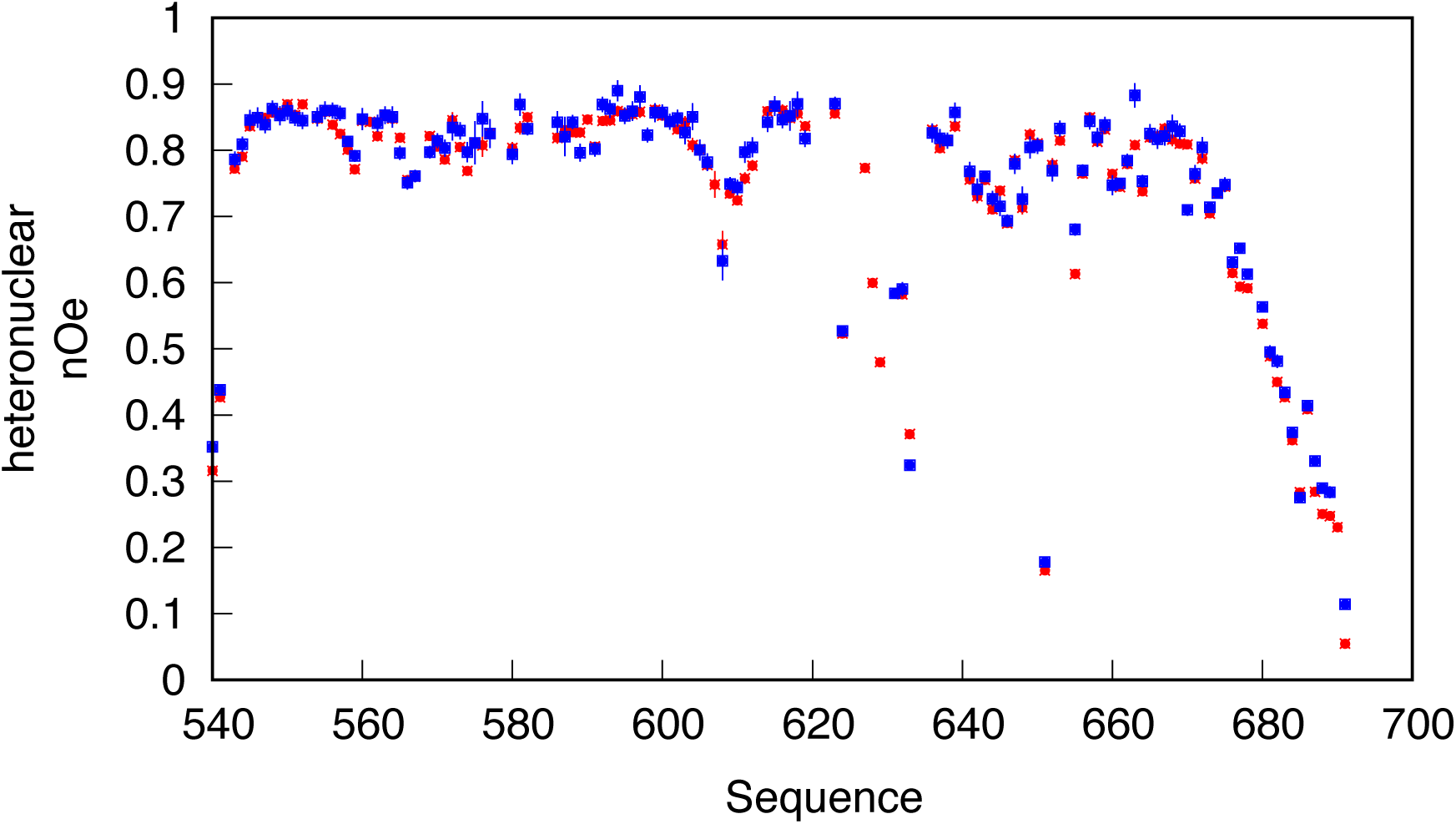
Heteronuclear nOe of 627K (red) and 627E (blue) HSQC. Data were recorded at 850MHz, 298 K.

**Figure S6.**
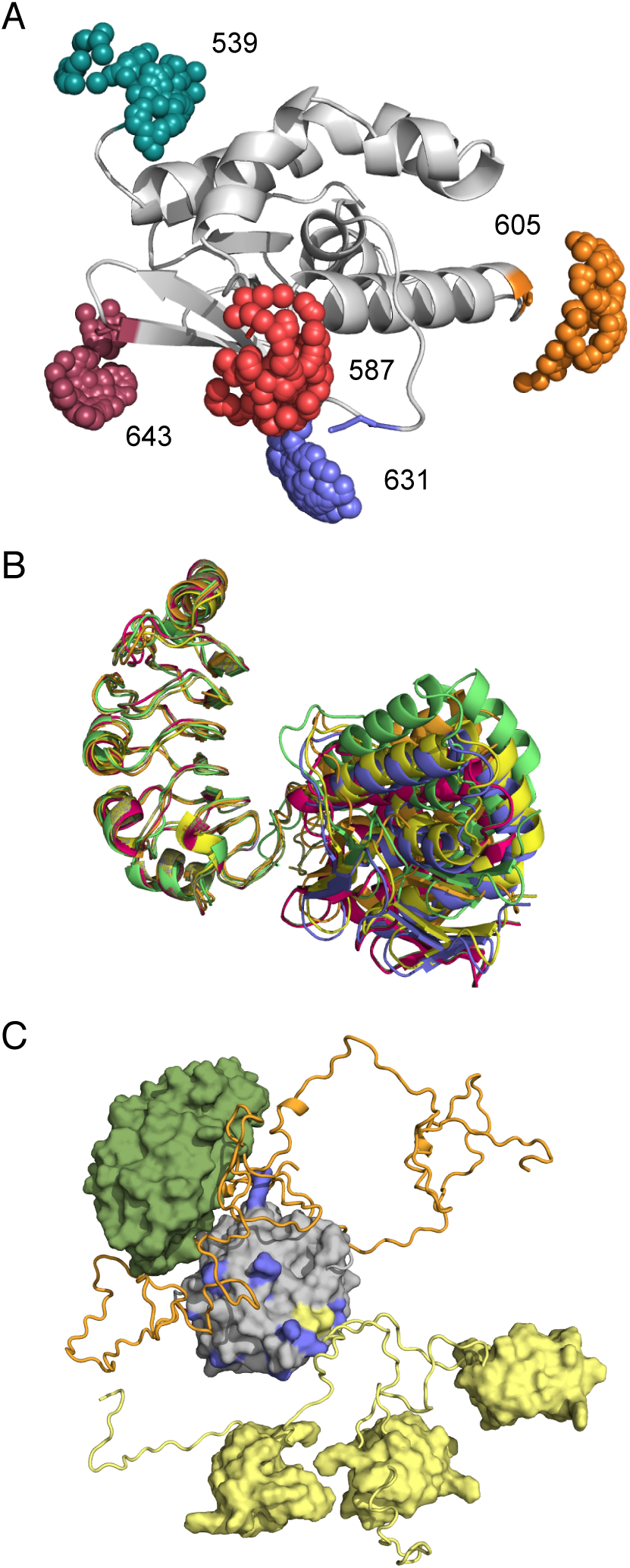
A - Representation of the sampling of TEMPO maleimide side-chain with respect to the back conformation of 627. Similar calculations were made for the three spin-label sites on the NLS domain (not shown). Rotamer-specific libraries were randomly sampled to place the nitroxide group in available conformations that respect steric clashes with the remainder of the molecule for each of the conformations sampled in the pool of conformations. B - Optimisation of the position of the folded domains of *h*ANP32A and 627-NLS(K). 2000 starting poses generated by the program Haddock^1^ were optimised with respect to experimental PREs measured on the folded domain. The 10 best fitting poses are shown. The same procedure was used for PREs measured on the *av*ANP32A: 627-NLS(E) complex, resulting in the same best-fitting structures. C - Representation of the generation of conformers present in the pool of 10000 conformers generated using the flexible-meccano algorithm. The statistical coil model was used to randomly sample the conformational space available to the linker region between the 627 (grey) and NLS (yellow) domains, the NLS peptide constituting the C-terminus of the 627-NLS domains, and the IDD domain of ANP32a (orange). The folded domain of ANP32a as shown in surface representation (green).

**Figure S7.**
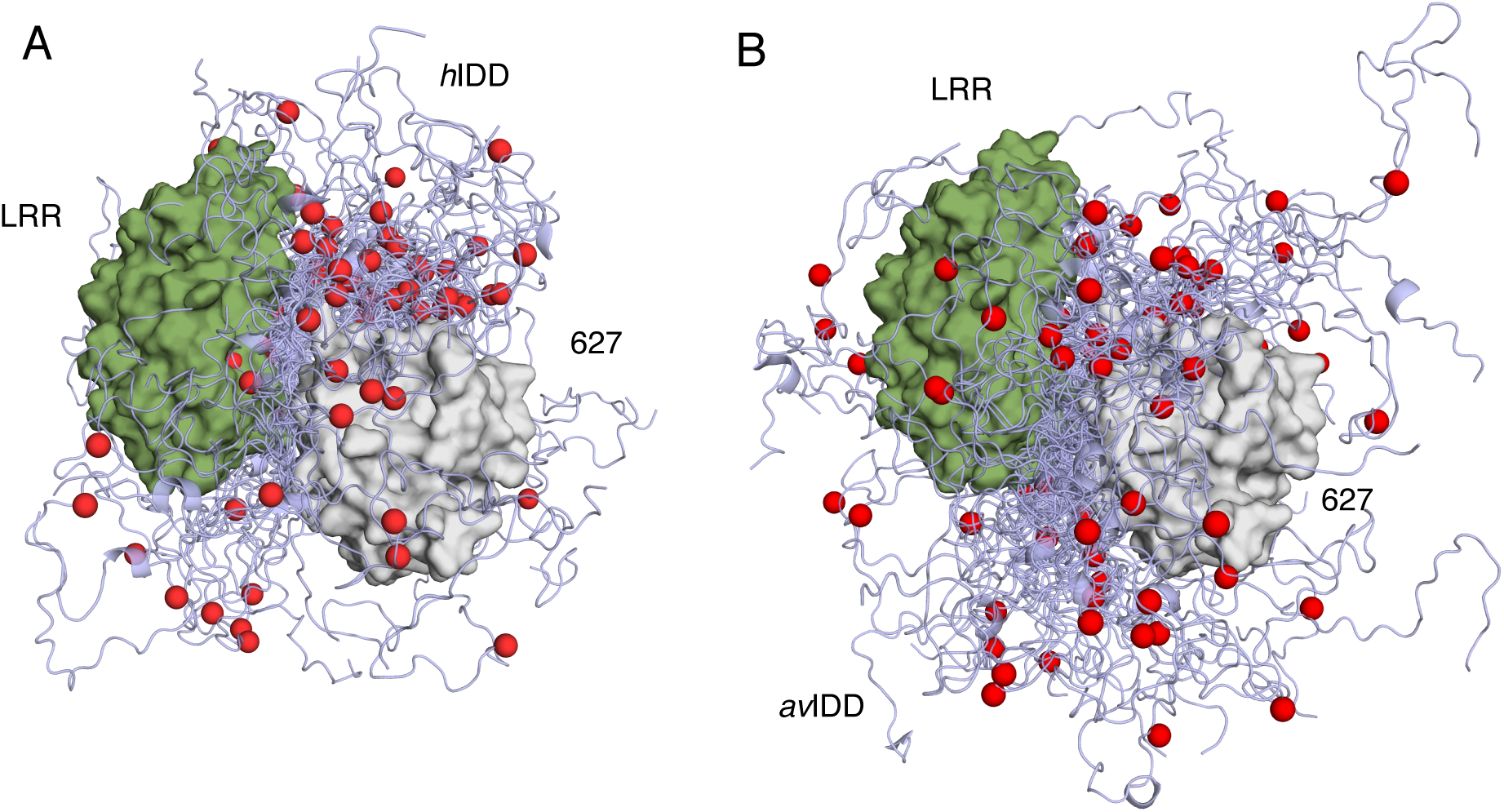
A. Representative ensemble of conformations describing the conformational space sampled by the *h*ANP32A:627-NLS(K) complex. The NLS domain has been removed for clarity, and the IDD truncated at residue 210. The position of residue 190 is indicated as a red sphere. B. Representative ensemble of conformations describing the conformational space sampled by the *av*ANP32A: 627-NLS(E) complex. Representation as in (B).

**Figure S8.**
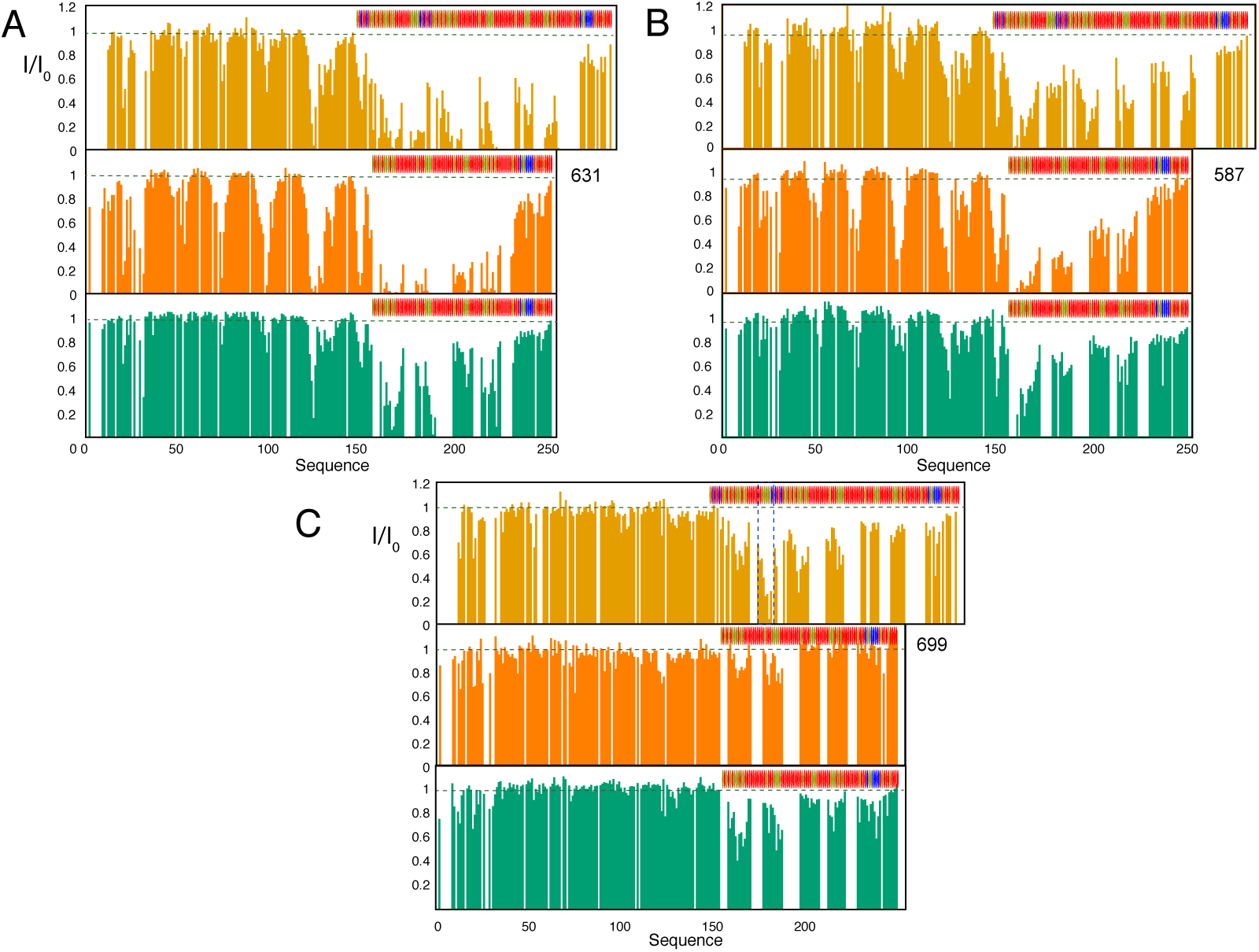
Comparison of intermolecular contacts in *h*ANP32A:627-NLS(K) (light orange), *av*ANP32A:627-NLS(E) (dark orange) and *h*ANP32A:627-NLS(E) (green). Three PREs are compared, (A) 631, (B) 587 and (C) 699. The comparison of PRE profiles due to the presence of a spin probe on residue 631 illustrates the reduced number and strength of contacts of the IDD with the 627 domain in the *h*ANP32A:627-NLS(E) complex compared to *h*ANP32A:627-NLS(K). 587 shows a similar effect. Finally comparison of PRE profiles due to the presence of a spin probe on residue 699 illustrates the lack of interaction with the linker-NLS region in the *h*ANP32A:627-NLS(E) compared to *av*ANP32A:627-NLS(E). The position of the hexapeptide region is identified with dotted vertical lines in C.

**Figure S9.**
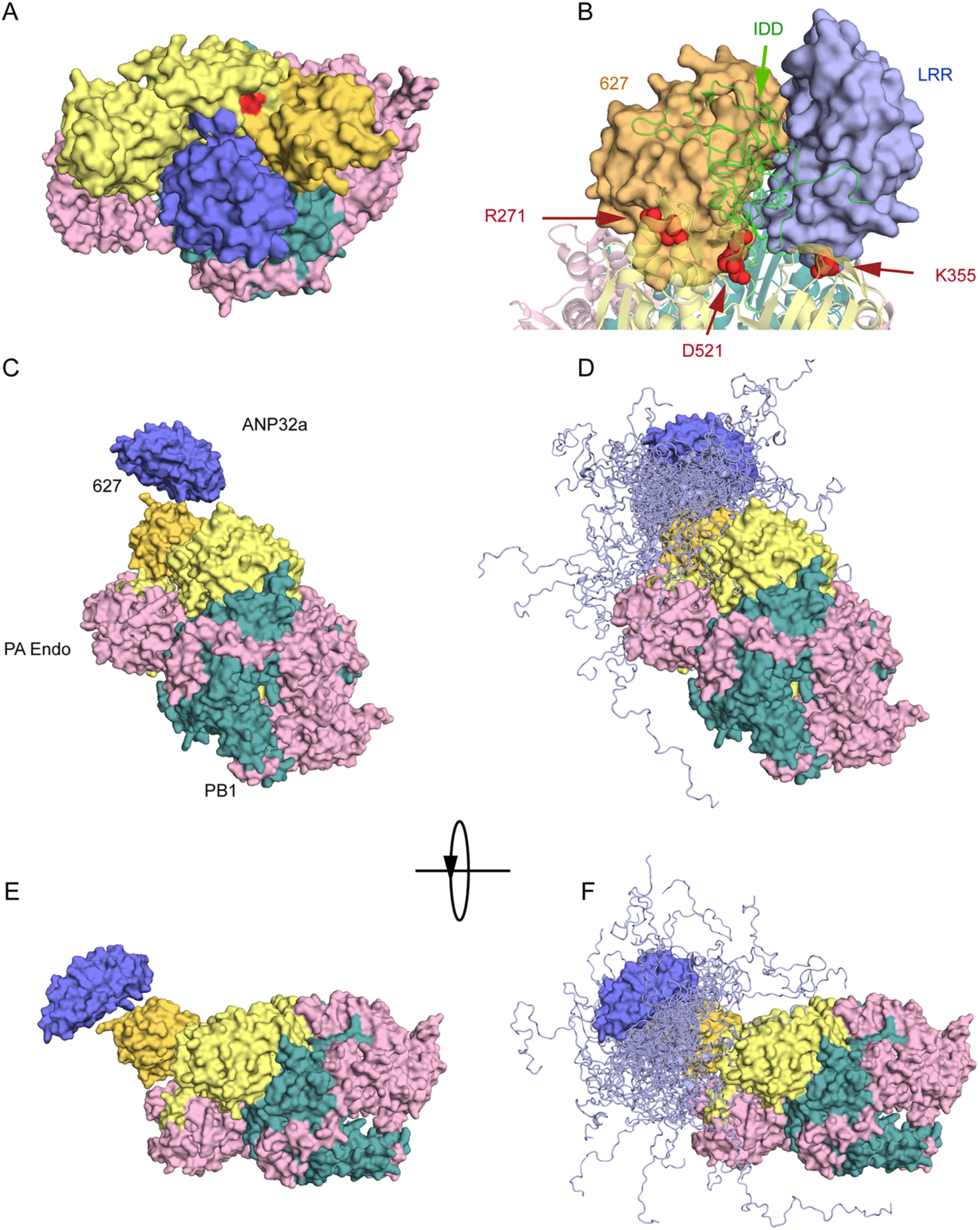
Binding pose of ANP32A with respect to the structure of bat Influenza B polymerase bound to *v*RNA promoter (4wsa).^2^ ANP32A LRR (blue surface) is positioned in a large pocket formed by 627 (yellow-orange) and cap-binding and mid domains of PB2 (yellow) and bordered by PB1 (dark-cyan). PB2 adaptive mutants R271, D521 and K355 all lie in the immediate vicinity of either 627 domain (R271), the interface between 627 and ANP32A LRR (D521) or ANP32A LRR (355). The first 25 amino acids of ten structures from the representative hANP32A:627-NLS(K) ensemble are shown to illustrate the beginning of the flexible IDD (green cartoon). C-F. Compatibility of binding mode determined in free solution in the context of the apo conformation of influenza A.^3^ C-D. The 627 domain was superimposed on the 627 domain of PB2 in the full length polymerase structure (6qnw). In this position, ANP32a folded domain can be adjacent to 627 (yellow-orange) on the surface of the polymerase. E-F. Conformational sampling of the IDD of *h*ANP32a, assuming the position of the folded domain of ANP32a shown in A and B. The linker and NLS domains are not shown for clarity and are assumed flexible.

## REFERENCES

1. Lozano, R. et al. Global and regional mortality from 235 causes of death for 20 age groups in 1990 and 2010: a systematic analysis for the Global Burden of Disease Study 2010. The Lancet 380, 2095–2128 (2012).

2. Nicholls, H. Pandemic Influenza: The Inside Story. PLOS Biol 4, e50 (2006).

3. Almond, J. W. A single gene determines the host range of influenza virus. Nature 270, 617–618 (1977).

4. Tarendeau, F. et al. Host determinant residue lysine 627 lies on the surface of a discrete, folded domain of influenza virus polymerase PB2 subunit. PLoS Pathog. 4, e1000136 (2008).

5. Mehle, A. & Doudna, J. A. Adaptive strategies of the influenza virus polymerase for replication in humans. Proc. Natl. Acad. Sci. U. S. A. 106, 21312–21316 (2009).

6. Gabriel, G., Czudai-Matwich, V. & Klenk, H.-D. Adaptive mutations in the H5N1 polymerase complex. Virus Res. 178, 53–62 (2013).

7. Tarendeau, F. et al. Structure and nuclear import function of the C-terminal domain of influenza virus polymerase PB2 subunit. Nat. Struct. Mol. Biol. 14, 229–233 (2007).

8. Subbarao, E. K., Kawaoka, Y. & Murphy, B. R. Rescue of an influenza A virus wild-type PB2 gene and a mutant derivative bearing a site-specific temperature-sensitive and attenuating mutation. J. Virol. 67, 7223–7228 (1993).

9. Massin, P., van der Werf, S. & Naffakh, N. Residue 627 of PB2 is a determinant of cold sensitivity in RNA replication of avian influenza viruses. J. Virol. 75, 5398–5404 (2001).

10. de Jong, M. D. et al. Fatal outcome of human influenza A (H5N1) is associated with high viral load and hypercytokinemia. Nat. Med. 12, 1203–1207 (2006).

11. Kirui, J., Bucci, M. D., Poole, D. S. & Mehle, A. Conserved features of the PB2 627 domain impact influenza virus polymerase function and replication. J. Virol. 88, 5977–5986 (2014).

12. Reilly, P. T., Yu, Y., Hamiche, A. & Wang, L. Cracking the ANP32 whips: important functions, unequal requirement, and hints at disease implications. BioEssays News Rev. Mol. Cell. Dev. Biol. 36, 1062–1071 (2014).

13. Long, J. S. et al. Species difference in ANP32A underlies influenza A virus polymerase host restriction. Nature 529, 101–104 (2016).

14. Sugiyama, K., Kawaguchi, A., Okuwaki, M. & Nagata, K. pp32 and APRIL are host cell-derived regulators of influenza virus RNA synthesis from cRNA. eLife 4, e08939 (2015).

15. Lowen, A. C. Virology: Host protein clips bird flu’s wings in mammals. Nature 529, 30–31 (2016).

16. Mehle, A. The Avian Influenza Virus Polymerase Brings ANP32A Home to Roost. Cell Host Microbe 19, 137–138 (2016).

17. Baker, S. F., Ledwith, M. P. & Mehle, A. Differential Splicing of ANP32A in Birds Alters Its Ability to Stimulate RNA Synthesis by Restricted Influenza Polymerase. Cell Rep. 24, 2581-2588.e4 (2018).

18. Long, J. S., Mistry, B., Haslam, S. M. & Barclay, W. S. Host and viral determinants of influenza A virus species specificity. Nat. Rev. Microbiol. 17, 67 (2019).

19. Staller, E. et al. ANP32 Proteins Are Essential for Influenza Virus Replication in Human Cells. J. Virol. 93, e00217–19 (2019).

20. Zhang, H. et al. Fundamental Contribution and Host Range Determination of ANP32A and ANP32B in Influenza A Virus Polymerase Activity. J. Virol. 93, (2019).

21. Baker, S. F. & Mehle, A. ANP32B, or not to be, that is the question for influenza virus. eLife 8, e48084 (2019).

22. Long, J. S. et al. Species specific differences in use of ANP32 proteins by influenza A virus. eLife 8, e45066 (2019).

23. Kuzuhara, T. et al. Structural basis of the influenza A virus RNA polymerase PB2 RNA-binding domain containing the pathogenicity-determinant lysine 627 residue. J. Biol. Chem. 284, 6855–6860 (2009).

24. Pflug, A., Guilligay, D., Reich, S. & Cusack, S. Structure of influenza A polymerase bound to the viral RNA promoter. Nature 516, 355–360 (2014).

25. Delaforge, E. et al. Large-Scale Conformational Dynamics Control H5N1 Influenza Polymerase PB2 Binding to Importin α. J. Am. Chem. Soc. 137, 15122–15134 (2015).

26. Hengrung, N. et al. Crystal structure of the RNA-dependent RNA polymerase from influenza C virus. Nature 527, 114–117 (2015).

27. Thierry, E. et al. Influenza Polymerase Can Adopt an Alternative Configuration Involving a Radical Repacking of PB2 Domains. Mol. Cell 61, 125–137 (2016).

28. Yamada, S. et al. Biological and structural characterization of a host-adapting amino acid in influenza virus. PLoS Pathog. 6, e1001034 (2010).

29. Salmon, L. et al. NMR Characterization of Long-Range Order in Intrinsically Disordered Proteins. J. Am. Chem. Soc. 132, 8407–8418 (2010).

30. Reich, S. et al. Structural insight into cap-snatching and RNA synthesis by influenza polymerase. Nature 516, 361–366 (2014).

31. Hayashi, T., Wills, S., Bussey, K. A. & Takimoto, T. Identification of Influenza A Virus PB2 Residues Involved in Enhanced Polymerase Activity and Virus Growth in Mammalian Cells at Low Temperatures. J. Virol. 89, 8042–8049 (2015).

32. Soh, Y. S., Moncla, L. H., Eguia, R., Bedford, T. & Bloom, J. D. Comprehensive mapping of adaptation of the avian influenza polymerase protein PB2 to humans. eLife 8, e45079 (2019).

33. Fan, H. et al. Structures of influenza A virus RNA polymerase offer insight into viral genome replication. Nature 573, 287–290 (2019).

34. Jensen, M. R., Zweckstetter, M., Huang, J. & Blackledge, M. Exploring free-energy landscapes of intrinsically disordered proteins at atomic resolution using NMR spectroscopy. Chem. Rev. 114, 6632–6660 (2014).

35. Milles, S. et al. Plasticity of an Ultrafast Interaction between Nucleoporins and Nuclear Transport Receptors. Cell 163, 734–745 (2015).

36. Borgia, A. et al. Extreme disorder in an ultrahigh-affinity protein complex. Nature 555, 61–66 (2018).

37. Fodor, E. & Te Velthuis, A. J. W. Structure and Function of the Influenza Virus Transcription and Replication Machinery. Cold Spring Harb. Perspect. Med. (2019) doi: 10.1101/cshperspect.a038398.

38. Büssow, K. et al. Structural genomics of human proteins--target selection and generation of a public catalogue of expression clones. Microb. Cell Factories 4, 21 (2005).

39. Braman, J., Papworth, C. & Greener, A. Site-directed mutagenesis using double-stranded plasmid DNA templates. Methods Mol. Biol. Clifton NJ 57, 31–44 (1996).

40. Delaglio, F. et al. NMRpipe - A Multidimensional Spectral Processing System Based On Unix Pipes. J. Biomol. NMR 6, 277–293 (1995).

41. Goddard, T. & Kneller, D. SPARKY 3. University of California, San Francisco.

42. Jung, Y.-S. & Zweckstetter, M. Mars -- robust automatic backbone assignment of proteins. J. Biomol. NMR 30, 11–23 (2004).

43. de Chiara, C., Kelly, G., Frenkiel, T. A. & Pastore, A. NMR assignment of the leucine-rich repeat domain of LANP/Anp32a. J. Biomol. NMR 38, 177 (2007).

44. Fossat, M. J. et al. High-Resolution Mapping of a Repeat Protein Folding Free Energy Landscape. Biophys. J. 111, 2368–2376 (2016).

45. Marsh, J. A., Singh, V. K., Jia, Z. & Forman-Kay, J. D. Sensitivity of secondary structure propensities to sequence differences between alpha- and gamma-synuclein: implications for fibrillation. Protein Sci. Publ. Protein Soc. 15, 2795–2804 (2006).

46. Lakomek, N.-A., Ying, J. & Bax, A. Measurement of 15N relaxation rates in perdeuterated proteins by TROSY-based methods. J. Biomol. NMR 53, 209–221 (2012).

47. Huang, J.-R. et al. Transient Electrostatic Interactions Dominate the Conformational Equilibrium Sampled by Multidomain Splicing Factor U2AF65: A Combined NMR and SAXS Study. J. Am. Chem. Soc. 136, 7068–7076 (2014).

48. Ozenne, V. et al. Flexible-meccano: a tool for the generation of explicit ensemble descriptions of intrinsically disordered proteins and their associated experimental observables. Bioinforma. Oxf. Engl. 28, 1463–1470 (2012).

49. Kubán, V. et al. Quantitative Conformational Analysis of Functionally Important Electrostatic Interactions in the Intrinsically Disordered Region of Delta Subunit of Bacterial RNA Polymerase. J. Am. Chem. Soc. 141, 16817–16828 (2019).

## REFERENCES

1. Dominguez, C., Boelens, R. & Bonvin, A. M. J. J. HADDOCK: a protein-protein docking approach based on biochemical or biophysical information. J. Am. Chem. Soc. 125, 1731–1737 (2003).

2. Reich, S. et al. Structural insight into cap-snatching and RNA synthesis by influenza polymerase. Nature 516, 361–366 (2014).

3. Fan, H. et al. Structures of influenza A virus RNA polymerase offer insight into viral genome replication. Nature 573, 287–290 (2019).

